# Demographic trade-offs decouple pollination services from plant population growth

**DOI:** 10.64898/2026.05.20.726591

**Authors:** Amy M. Iler, Paul J. CaraDonna, William K. Petry

**Affiliations:** Negaunee Institute for Plant Conservation Science and Action, Chicago Botanic Garden, 1000 Lake Cook Road, Glencoe, IL 60022, USA; Plant Biology and Conservation, Northwestern University, 633 Clark St, Evanston, IL 60208, USA; Rocky Mountain Biological Laboratory, 8000 County Road 317, Crested Butte, CO 81224, USA; Department of Plant and Microbial Biology, North Carolina State University, Raleigh, NC 27695, USA

**Keywords:** cost of reproduction, demographic buffering, experimental demography, integral projection model, pollinator decline, resource reallocation, trade-off

## Abstract

Most plants require animal pollination to reproduce, prompting concern that pollinator declines immediately threaten plant populations. This concern is warranted if pollinator-mediated seed losses cause declines in plant population growth rates (λ). However, demographic trade-offs might reduce the risk of population decline if seed loss improves performance elsewhere in the life cycle. We conducted a multi-year pollination manipulation on four species and measured how demographic vital rates and λ responded. Seed responses did not predict net changes in λ. Reduced pollination decreased seed production, but only caused a net decrease in λ in one species; in the others, improved survival buffered λ. Increased pollination boosted seed production, but at a cost to survival that caused a net reduction in λ in three species. Our results highlight the importance of demographic trade-offs for understanding the impacts of pollinator declines on plant biodiversity and, more broadly, the population-level impacts of changing mutualisms.

## Introduction

The effects of mutualistic interactions on plant populations are poorly understood (Ehrlén *et al*. 2016; Morris *et al*. 2020), and this gap in our understanding is especially relevant in light of widespread evidence of pollinator declines (Biesmeijer *et al*. 2006; Burkle *et al*. 2013; Goulson *et al*. 2015; Potts *et al*. 2016). Declines in pollination services are expected to cause plant population declines and co-extinctions because most plant species reproduce with the aid of animal pollinators (Ollerton *et al*. 2011; Rodger *et al*. 2021). Indeed, there is observational evidence that plant species are already declining in response to pollinator declines (Anderson *et al*. 2011; Biesmeijer *et al*. 2006). At the same time, the expectation of pollinator-mediated plant decline rests on a rarely tested tenet of pollination biology—that changes in pollination have strong effects on plant fitness (Lundgren *et al*. 2015; Price *et al*. 2008; Thomson 2025). Pollinator-mediated changes in seed production are largely studied in isolation from plant population dynamics. Although many pollination studies mechanistically measure fitness components like seed production and seed siring, these fitness components are typically isolated from the rest of the plant life cycle (Lundgren *et al*. 2015; Price *et al*. 2008). This is especially true in the context of declines in pollination services.

In contrast to expectations and evidence from pollination biology, demographic life-history theory suggests that many plant species should be buffered from the negative consequences of pollinator declines because of trade-offs and plant life histories. Demographic trade-offs are common because plants have a finite pool of resources, and they occur when alterations to one vital rate cause compensatory changes in other vital rates (Stearns 1992). For example, if *declines* in pollination reduce seed production, plants might fully or partially offset effects of seed losses on population growth by reallocating reproductive resources to future survival and vegetative growth. Conversely, if *increases* in pollination increase seed production, this greater cost of reproduction might negatively affect other vital rates (Ehrlén & Eriksson 1995; Obeso 2002). Such trade-offs between reproduction and other vital rates are possible for the vast majority of plant species, which have an iteroparous, perennial life-history strategy that distributes reproductive events across multiple years (Friedman 2020; Poppenwimer *et al*. 2023; Young & Augspurger 1991). In particular, populations of iteroparous perennials might be intrinsically buffered against reproductive losses under pollinator declines because their population growth rates tend to be proportionately less responsive to perturbations in reproduction and more responsive to perturbations in somatic growth and survival (Franco & Silvertown 2004; Pfister 1998; see *Supporting Information, section 1* for a hypothetical example). Therefore, even small trade-offs of reproduction on growth and survival could have large effects on population growth. Although these expectations from demographic life-history theory are well-known to demographers, they are underappreciated in the broader ecological literature in which changes in reproduction are frequently assumed to have strong effects on populations and communities, regardless of life history (*reviewed by* Crone 2001; Iler *et al*. 2021).

Previous experimental work linking changes in pollination to changes in plant population growth has focused on measuring the effects of *increased* pollination to determine whether plant populations are limited by pollination (Baer & Maron 2018; Ehrlén & Eriksson 1995; Knight 2004; Price *et al*. 2008). The consequences of experimental *decreases* in pollination for plant population growth are virtually unknown, thereby largely obscuring the consequences of pollinator declines for most plant biodiversity on the planet (*i.e.,* iteroparous perennials; see Johnson *et al*. 2022 for a study on annuals). Because of demographic trade-offs, population-level responses to decreased pollination are unlikely to simply mirror the effects of increased pollination in the opposite direction, highlighting the importance of studying both increases and decreases in pollination in the same species.

A whole life-cycle approach will help to resolve the conflicting expectations between pollination biology and life-history theory about how changes in pollination might affect population outcomes for plants. We therefore conducted an experimental demography study wherein we manipulated pollination (increased, decreased, and ambient pollination) and measured responses across the entire life cycle of four herbaceous plant species across multiple years in the Colorado Rocky Mountains, USA. We ask: (1) What are the net effects of both decreases and increases in pollination on the population growth rates of iteroparous, perennial plant species (quantified by, λ, the finite rate of increase or mean fitness)? (2) How well do changes in seed production predict the net treatment effect on λ? (3) How is λ impacted by non-reproductive vital rate responses to pollination treatment through demographic trade-offs? The simplest hypothesis is that decreased pollination will reduce seed production, which will in turn reduce λ (and vice versa). However, we expect that responses of λ to the pollination treatments will ultimately depend on: (i) the magnitude of gains and losses in seed production, (ii) the strength of trade-offs among vital rates, and (iii) how responsive λ is to perturbations to each vital rate.

## Materials and Methods

### Study system & species

The four herbaceous, iteroparous perennial plant species in this study were *Delphinium nuttallianum* (Ranunculaceae)*, Hydrophyllum fendleri* (Hydrophyllaceae), *Potentilla pulcherrima* (Rosaceae), and *Erigeron speciosus* (Asteraceae), hereafter referred to by genus (Table 1; Fig. S3). All species are relatively common in the Colorado Rocky Mountains, reproduce through seed, but differ in lifespan, primary pollinator species, and ability to produce seed via self-pollination (Table 1; Fig. S3; detailed natural history information is available in *Supporting Information, 2a*). We worked near the Rocky Mountain Biological Laboratory (RMBL; 2900 m asl; 38.968° N, 106.998° W) from 2017 through 2022, amassing five years of demographic census data for *Hydrophyllum* and four years of data for the other three species (2018–2021 for *Potentilla* and *Erigeron*; 2019–2022 for *Delpshinium*). The landscape around the RMBL is a patchwork of montane meadows dominated by herbaceous perennial plants, intermixed with aspen and conifer forest, with herbaceous perennials also dominating the understory.

**Table 1.**
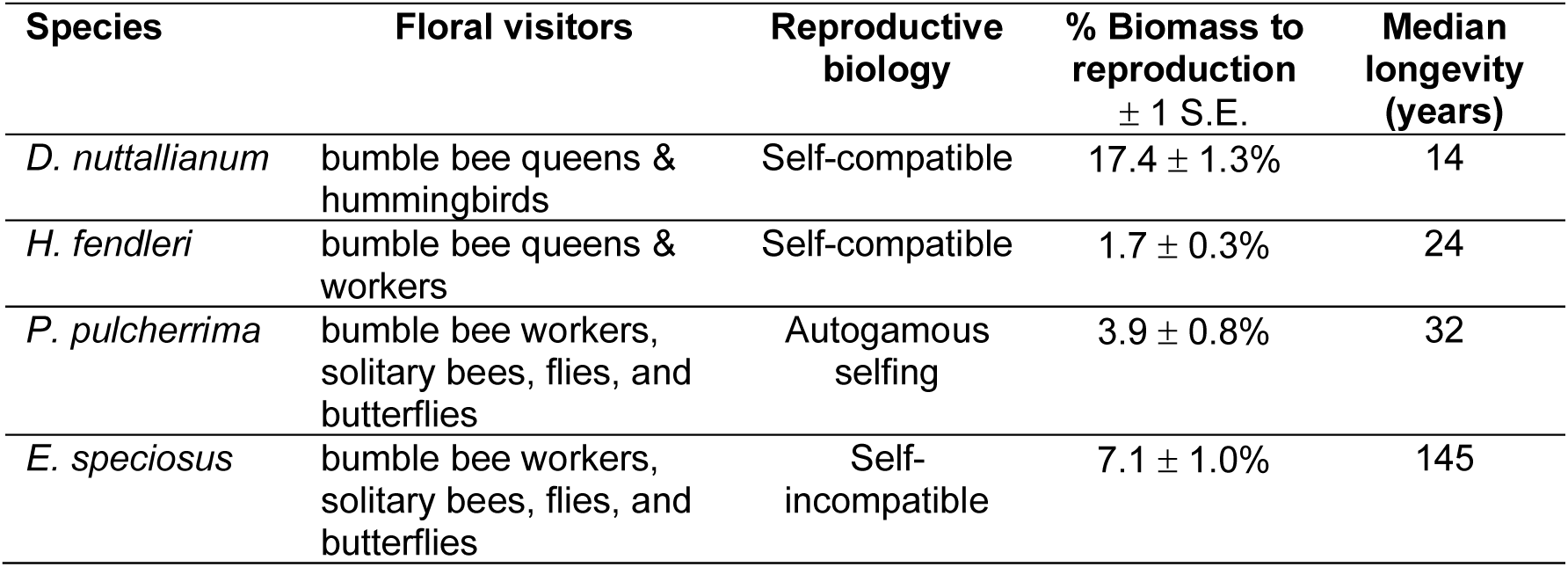
Pollination biology and life history information for the four study species: *Delphinium nuttallianum*, *Hydrophyllum fendleri*, *Potentilla pulcherrima*, and *Erigeron speciosus.* Methods for quantifying the percent of total biomass allocated to reproductive effort are detailed in *Supporting Information, 2d*. Reproductive biomass is represented by flowering stalks with flowers for *P. pulcherrima* and *E. speciosus*, flowering stalks with flowers and some fruits for *H. fendleri*, and only the flowering stalk, with no flowers or fruits, for *D. nuttallianum* (for a conservative estimate of % biomass allocated to reproductive effort in *D. nuttallianum*). Longevity was calculated as the 99^th^ percentile of lifespan from IPMs: specifically, we made 1,000 IPM kernels from vital rates drawn from the posterior distribution, simulated a cohort, and calculated the median number of years it took for 99% of them to die.

| Species | Floral visitors | Reproductive biology | % Biomass to reproduction<br>$\pm 1$ S.E. | Median longevity (years) |
| --- | --- | --- | --- | --- |
| *D. nuttallianum* | bumble bee queens & hummingbirds | Self-compatible | $17.4 \pm 1.3\%$ | 14 |
| *H. fendleri* | bumble bee queens & workers | Self-compatible | $1.7 \pm 0.3\%$ | 24 |
| *P. pulcherrima* | bumble bee workers, solitary bees, flies, and butterflies | Autogamous selfing | $3.9 \pm 0.8\%$ | 32 |
| *E. speciosus* | bumble bee workers, solitary bees, flies, and butterflies | Self-incompatible | $7.1 \pm 1.0\%$ | 145 |

### Experimental Design

We first designated a series of study plots that sampled a single population of each species. We selected plot locations by measuring the length and width of our study populations and drawing random coordinate pairs contained within the population. Each plot was 1 m × 10 m, was divided in half lengthwise, and was further divided into four treatment areas per half plot, each 0.5 m × 2.5 m. Within each half plot, we randomly assigned each area to one of four pollination treatments: unmanipulated control (ambient pollination), decreased pollination, increased pollination, or variable pollination (in which pollination was decreased in only one of the study years). Plants in the variable pollination treatment were excluded from this study. All individuals within the treatment area received the respective treatment.

To decrease pollination, we aimed to enclose half of the flowers on an individual plant in a pollinator exclusion bag that bees and other flower visitors could not penetrate (Kearns & Inouye 1993). If there were an odd number of flowers on a plant, we erred on the side of bagging more flowers (*e.g.,* if a plant had three flowers, two were bagged; if a plant had five flowers, 3 were bagged).

To increase pollination, we added supplemental outcross pollen to stigmas during the morning, when anthers contained most pollen, typically between 0800 and 1000 hours. We augmented natural pollen delivery to all receptive stigmas on the flowers of an individual plant by adding outcross pollen by hand three times per week during the flowering period of each species. Conspecific pollen donor plants were selected from the same population, but outside the plots and at least 5 m away from focal plants. We removed mature anthers from donor plants, then gently rubbed them across stigmas of experimental plants until stigmas were visibly coated with pollen (Ashman *et al*. 2004; Kearns & Inouye 1993). The individual flowers of *Erigeron* are too small to hand pollinate individually; therefore, we gently rubbed entire donor capitula (flower heads) containing visible pollen against the capitula of experimental plants with visible stigmas.

Across all plots for each species, we tagged at least 250 individuals per treatment with a unique identification number. We then we collected individual-level demographic data from these plants annually for at least four years. Plant density varied among species and determined the number of plots necessary for including at least 250 individuals per treatment: four plots for *Potentilla*, six plots for *Delphinium* and *Erigeron,* and 16 plots for *Hydrophyllum*. Each year, we measured the following demographic data from all tagged individuals in our study plots: survival (yes/no), plant size (metric varied by species, details below), reproductive status (flowering: yes/no), reproductive effort (# flowers), and reproductive output (# seeds). We chose the size metric for each species based on its skill in predicting the vital rates (see below) from a pool of several candidate metrics tailored to the species’ morphology (Gamelon *et al*. 2021). For *Delphinium*, we used an approximation of leaf area multiplied by leaf number [the formula for an ellipse was used for leaf area: (leaf length × 0.5) (leaf width × 0.5) × π]; for *Hydrophyllum* we used the number of leaves; for *Potentilla* we used length of the longest leaf; and for *Erigeron* we used the height of the tallest stalk. We collected fruits as they matured, dried them, and counted seeds in the laboratory. We counted all seeds from reproductive individuals, except for *Erigeron*, where it was not feasible to individually count the large number of seeds per individual. Instead, we counted the number of seeds per capitulum from a sample of capitula per plant and multiplied that by the number of capitulae to estimate plant-level seed production. In total across all years and species, we measured the fates of >4,000 individual plants across >12,500 annual demographic transitions (*i.e.,* individual vital rates from one year to the next).

To determine whether the seed-to-seedling transition was density dependent, we conducted seed addition experiments adjacent to the demography plots. We sowed seeds into areas containing adults of each focal species and bare ground. We created sowing density treatments ranging from no seeds (control) up to 10× the densest seed rain observed in the increased pollination treatment, with intervening densities evenly spaced on a log scale (*following* Price *et al*. 2008; *Supporting information, 2b*). Each density treatment was replicated three times for a total of 21 plots for species with seven sowing densities and 18 plots for species with six sowing densities (*Supporting Information*, *2b*.) All germination plots were circular and included a 1 m buffer around each plot from which fruits were removed to reduce natural seed rain into the plots (*following* Price *et al*. 2008). The year after sowing, we counted the number of seedlings in each plot and regressed the number of seedlings against the density of seeds sown for each species to look for evidence of density-dependent recruitment (using GLMs with a negative binomial distribution to account for overdispersion, *MASS* package, Venables & Ripley 2002).

We did not find evidence for density dependence in the seed to seedling transition in any of the four species and therefore did not incorporate these dynamics into our population models. In *Delphinium* and *Hydrophyllum*, sowing density did not significantly affect the number of seedlings that emerged in the following year (*Delphinium: z* = -0.52, *p* = 0.60; *Hydrophyllum*: *z* = -0.24, *p* = 0.81, *n* = 18 plots for each). Consistent with the incredibly low number of seedlings that we observed in the field for *Potentilla* and *Erigeron,* no *Potentilla* seedlings emerged in our seed germination experiment, and only nine *Erigeron* seedlings emerged (out of 1800 seeds sown), for a mean germination rate of 0.5%. Germination rates this low, despite varying sowing densities tenfold, suggest that any biological impact of density dependence at this stage is negligible.

*Delphinium nuttallianum* does not have a seed bank in our study area (Waser & Price 1985), but the seed bank status of our other three study species was unknown, so we also conducted seed bank experiments and incorporated them into our population models as necessary (only for *Hydrophyllum*; *Supporting Information, 2c*).

### Demographic Models & Analyses

All analyses were performed in R v 4.5.0 (R Core Development Team, 2025). We used package *ipmr* to construct a deterministic, density-independent Integral Projection Model (IPM) tailored to the life history of each species (Easterling *et al*. 2000; Levin *et al*. 2021) (details of model construction are in *Supporting Information*, *2e*.) Briefly, we parameterized the IPMs with demographic vital rate functions estimated from a Bayesian multivariate, multilevel model fit to the field data under each pollination treatment. Each vital rate response for established plants included an intercept and effects of plant size, pollination treatment, and the size × treatment interaction as population-level (“fixed effect”) predictors. We fit group-level (“random effect”) intercepts for plot and year and allowed information to be shared among the vital rate responses through correlations between these intercepts. We estimated germination using data from seed addition plots, and the model fit a population-level intercept that accounted for natural seed rain and slope for the number of seeds. We used a single population-level intercept to estimate the seedling size distribution, omitting effects of treatment, plot, or year. We discretized the IPM projection kernel into a 400×400 matrix using the midpoint rule. We appended a discrete single-age class seedbank stage for *Hydropyllum*, the only species for which seed survival was observed. We projected the IPM for each species to equilibrium under each treatment using 1,000 parameter sets drawn from joint posterior distribution of the vital rate model. From these projections, we extracted the posterior distribution of λ for each species-treatment combination (Q1).

To understand whether changes in seed production predicted changes in λ (Q2), we calculated the percent change in both variables for each species and treatment comparison (reduced vs. control and increased vs. control for 4 species = 8 total values). For seed production, we used the percent change of mean values of seeds per plant across all study years. We summarized the posterior distribution of λ using the mean for each species and treatment. We then used a linear regression model with percent change in seeds as the predictor and percent change in λ as the response.

Because multiple vital rates could respond to pollination treatment, and each could have a different impact on λ, we used Life Table Response Experiments (LTREs) to decompose the net treatment effect on λ into the contributions from each underlying vital rate response (Q3; *Supporting Information, 2e*). Here too, we used the joint posterior distribution of vital rate parameters to appropriately propagate uncertainty from the vital rates to the LTRE.

Finally, we conducted an elasticity analyses under the control treatment to characterize the proportional change in λ in response to perturbations in different components of the life cycle (De Kroon *et al*. 1986), using the elasticity function in the *popbio* package (Stubben & Milligan 2007). We partitioned the kernel elasticities into additive components describing persistence (growth and survival) and reproduction, then summed the kernel elasticities within each partition (*Supporting Information, 2f*). We propagated the uncertainty in the IPM vital rate parameters by repeating this elasticity partitioning for 1,000 parameter sets drawn from the joint posterior distribution of the vital rate model, then calculated the median of the resulting proportions. This allowed us to further understand why vital rate trade-offs affect λ, in the context of life history.

Although evidentiary cut-offs in any statistical framework are arbitrary, we chose 90% and 75% Bayesian credible intervals as thresholds for strong and moderate evidence of pollination treatment effects, respectively, to facilitate qualitative discussion of results. We present the full posteriors in figures and in our archived data for further interpretation at other thresholds.

Like most studies, our demographic analyses assume that each population was near the equilibrium plant size distribution and population density prior to the start of our experiment. Because our study species are long-lived (Table 1), we needed to test whether our pollination treatments moved the treatment equilibrium far from the control, which could invalidate our LTRE analysis. We compared the equilibrium size distributions for each treatment with the control, and all remained close to the control in all species (Fig. S4). Only one species-treatment combination showed evidence of a change in the stable size distribution relative to the control; the magnitude of this difference was small and unlikely to affect the conclusions of our study (Fig. S4). Because density varies naturally across our sampling areas, it is implicitly built into our models but is not modeled explicitly. Our experiment focused on density-dependence at the recruitment stage because it is the stage at which we expected pollination treatment to have the strongest density-dependent effects, via changes in seed rain. Finally, we assumed that our study years captured the mean background environmental conditions that our populations experience, and our population models were designed to compare population growth rate responses to chronic pollination changes. The 8–10 years of data required to model stochastic variation in pollination or year-to-year environmental conditions was beyond the scope of this study (Caswell 2001; Doak *et al*. 2005; Ellner *et al*. 2016).

## Results

The pollination treatments affected seed set as expected (Table 2). All species experienced declines in the number of seeds/flower in response to decreased pollination (ranging from 15.3% – 63.4% reductions in mean seeds/flower per plant; Table 2). To a lesser extent, increased pollination increased seeds/flower in all four species (ranging from 2.8% – 17.7% increases; Table 2). We calculated treatment effect sizes using seeds/flower (vs. seeds/plant) because flower-level seed production is typically independent of plant size and therefore any potential trade-offs with size.

**Table 2.** Pollination treatments affect seed set in the expected direction. Percent change in the mean number of seeds per flower is shown, with the control as the reference value. Values are calculated from raw data: means for seeds per flower per individual across all four years of the experiment. The number of seeds per flower isolates the effect of the pollination treatments because this response typically occurs independent of plant size and therefore independent of potential compensation in other vital rates, due to costs of reproduction (as opposed to plant-level seed production, which would include effects of compensation, thereby confounding the direct effect of the pollination treatment on seed production). For *E. speciosus*, data are seeds per capitula or flowerhead, as each capitula contains numerous individual flowers.

| Species | Treatment comparison | % change in seeds per flower |
| --- | --- | --- |
| *D. nuttallianum* | decreased vs. control | -63.4 |
|  | increased vs. control | +2.8 |
| *H. fendleri* | decreased vs. control | -52.6 |
|  | increased vs. control | +11.8 |
| *P. pulcherrima* | decreased vs. control | -15.3 |
|  | increased vs. control | +7.3 |
| *E. speciosus* | decreased vs. control | -56.6 |
|  | increased vs. control | +17.7 |

Over the study period, mean λ values in the control treatments were 1.03 for *D. nuttallianum* (3% annual increase), 0.89 for *H. fendleri* (11% annual decrease), 0.94 for *P. pulcherrima* (6% annual decrease), and 1.20 for *E. speciosus* (20% annual increase). Changes in seed production failed to predict changes in λ (seeds per plant: *R^2^* = 0.07; *F^1,6^* = 0.44; *p* = 0.53, Fig. 1; seeds per flower: *R^2^* = 0.03, *F_1,6_* = 0.21, *p* = 0.66; *SI Appendix,* Table S1). Consistent with this, the net effects of the pollination treatments on λ varied in direction and magnitude, in large part due to trade-offs among vital rates (Fig. 2; Fig. S5).

**Fig. 1.**
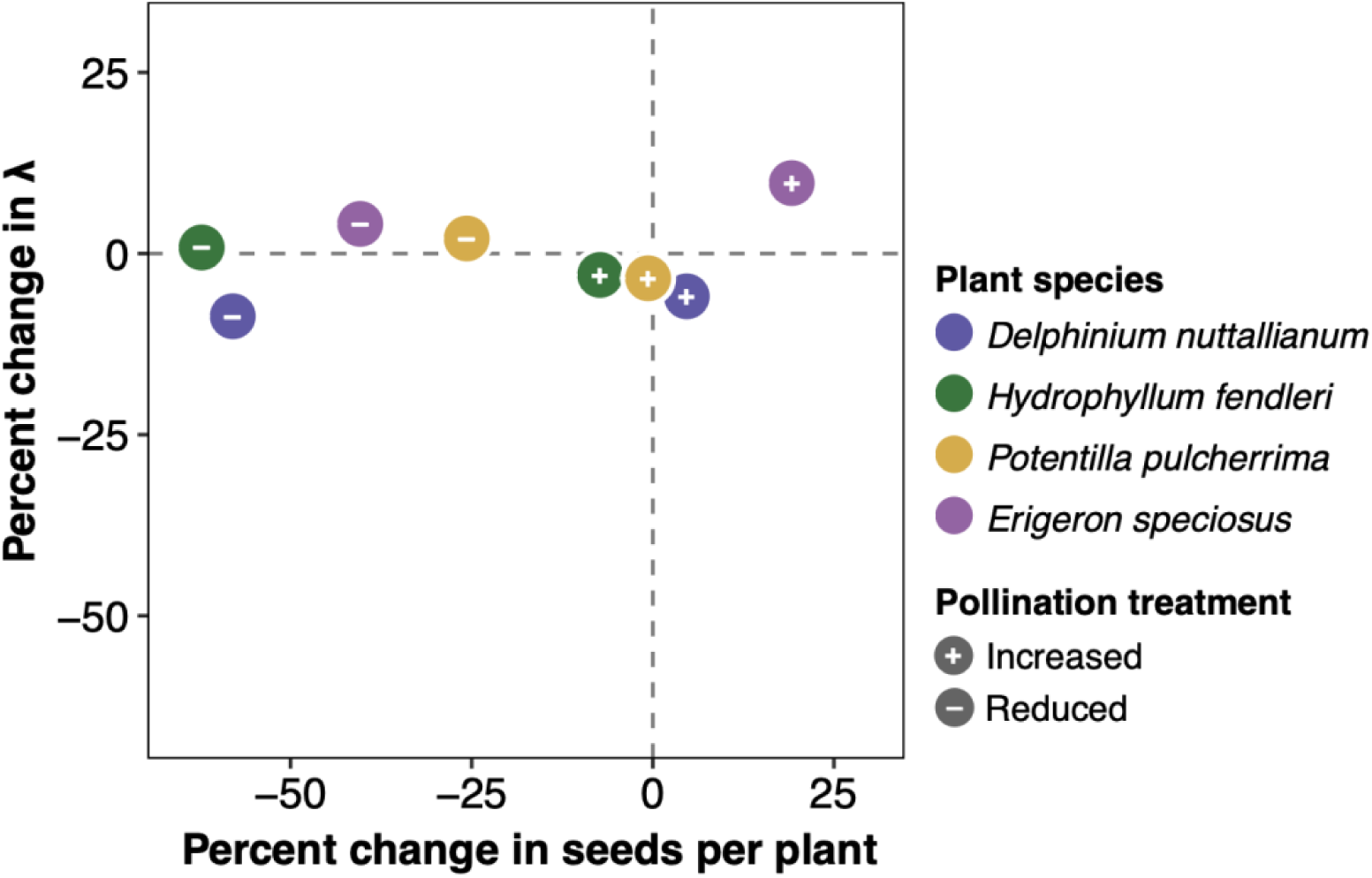
Changes in seed production in response to pollination treatments do not predict changes in λ. Each dot represents the magnitude of change in mean plant-level seed production vs. the magnitude of change in mean λ for a single plant species, in response to two pollination treatments: decreased (minus signs) and increased (plus signs) pollination. The magnitude of change is calculated as the percent change of mean values relative to the control for each species and treatment. Points above the horizontal dashed line are populations for which λ is projected to increase, and vice versa. Although flower-level seed production best represents treatment effects (Table 2), here we focused on the ability of change in plant-level seed production to predict change in λ, because in general, ecologists are most likely to use individual-level metrics to predict population-level responses. Regardless, change in seeds per flower explains even less variation in change in λ.

**Fig. 2.**
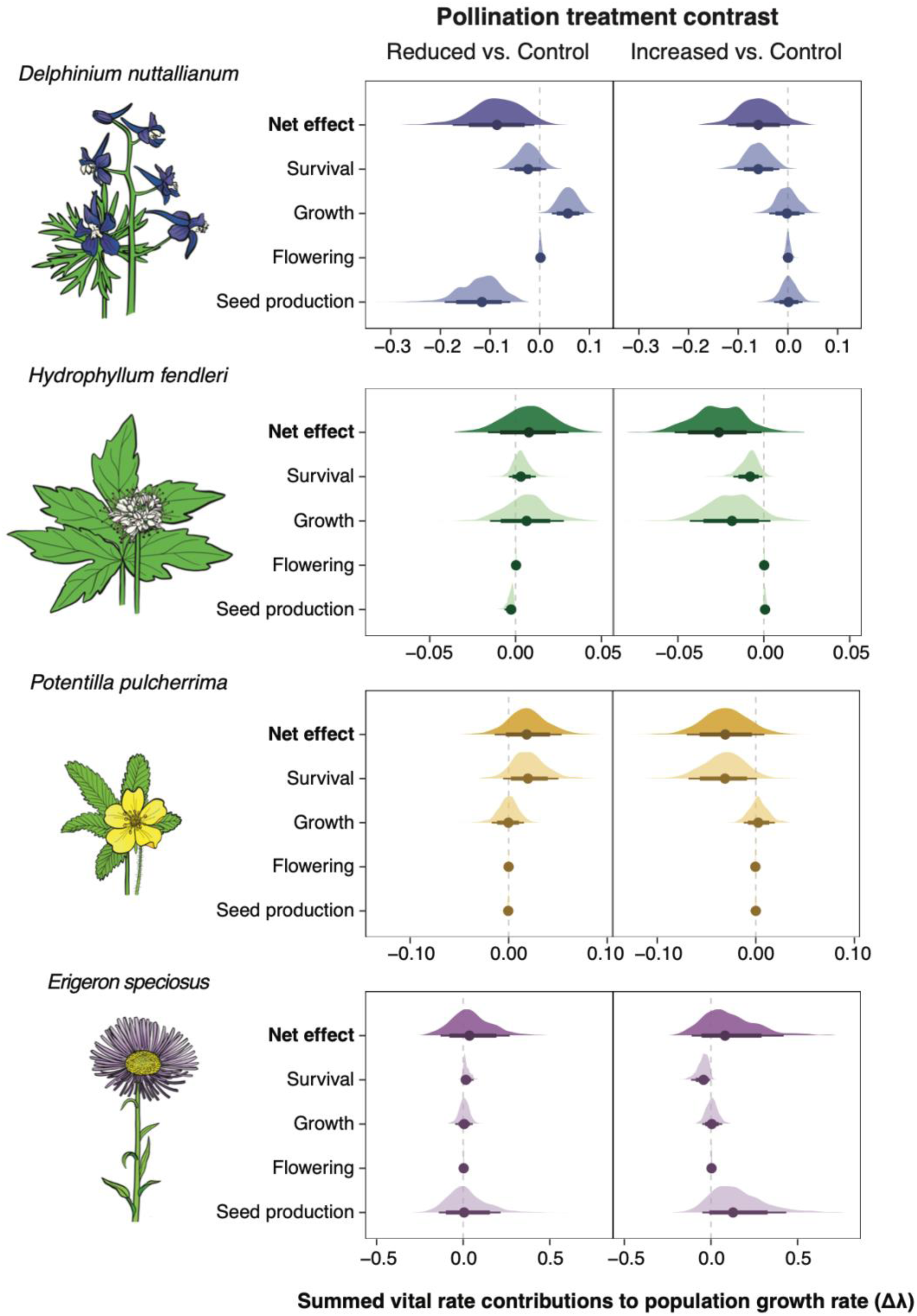
Changes in pollination affect λ directly, via changes to seed production, and indirectly through survival and vegetative growth that compensate for changes in seed production. Results are from a one-way fixed Life Table Response Experiment (LTRE), which determines how the change in population growth rate, Δλ, arose from the differences in the underlying vital rate parameters between the treatments and control. The vertical dashed grey line represents the control treatment. The net effect represents the net change in λ and is the summed contribution of each vital rate. The net effect is then broken down into contributions for each vital rate. Shaded regions show the full posterior distributions, thin lines are 90% credible intervals, thick lines are 75% credible intervals, and dots are medians. Note that the contribution of seed production can be near zero, not because the pollination treatments failed to affect seed production (see Fig. 1) but instead because changes in seed production do not have strong effects on λ. Plant illustrations by Julia Johnson (Life Science Studios).

Decreased pollination caused a net decrease in λ only in *Delphinium*, and the LTRE analysis revealed that this was largely due to a negative contribution from the decrease in seed production (Fig. 2). There was evidence of partial compensation from increases in vegetative growth, but this was insufficient to offset the negative effects of seed production on λ (Fig. 2).

In contrast to *Delphinium*, we found that λ was strongly buffered against the observed levels of seed loss under the decreased pollination treatment in the other three species (*Hydrophyllum, Potentilla*, and *Erigeron*), for which there was no net change in λ compared to the controls (*i.e.,* 75% credible interval overlaps 0; Fig. 2). Despite no net change in λ, the LTRE revealed vital rate contributions to changes in λ in both positive and negative directions. In *Hydrophyllum*, reduced seed production alone would have caused a net decrease in λ, but simultaneous positive contributions from survival and growth buffered the negative effects of seed production (Fig. 2; Fig. S5). In *Potentilla* and *Erigeron,* decreased pollination had almost no effect on λ via seed production, despite the reduction in seed production. Nonetheless, there was still evidence for trade-offs with survival that boosted λ (Fig. 2; Fig. S5).

In the increased pollination treatment, higher seed production consistently reduced survival, resulting in negative LTRE contributions of survival to λ in all four species (Fig. 2). In three species, the diversion of resources away from survival (and sometimes growth) was strong enough to result in a net decrease in λ relative to controls (strong evidence in *Hydrophyllum,* moderate evidence in *Delphinium* and *Potentilla*; Fig. 2).

*Erigeron* was the only case in which we found any evidence of a trend toward a net increase in λ under increased pollination (Fig. 2). Like the other three species, *Erigeron* experienced a cost of reproduction in terms of survival, which contributed negatively to λ (Fig. 2). However, the positive contribution of increased seed production offset this cost (Fig. 2).

For all species, the elasticity analysis revealed higher elasticities of λ to perturbations in survival and vegetative growth rather than reproduction (Fig. 3). *Delphinium* and *Erigeron* showed moderate elasticities of λ to reproduction (Fig. 3), consistent with a greater capacity for λ to respond to changes in pollination.

**Fig. 3.**
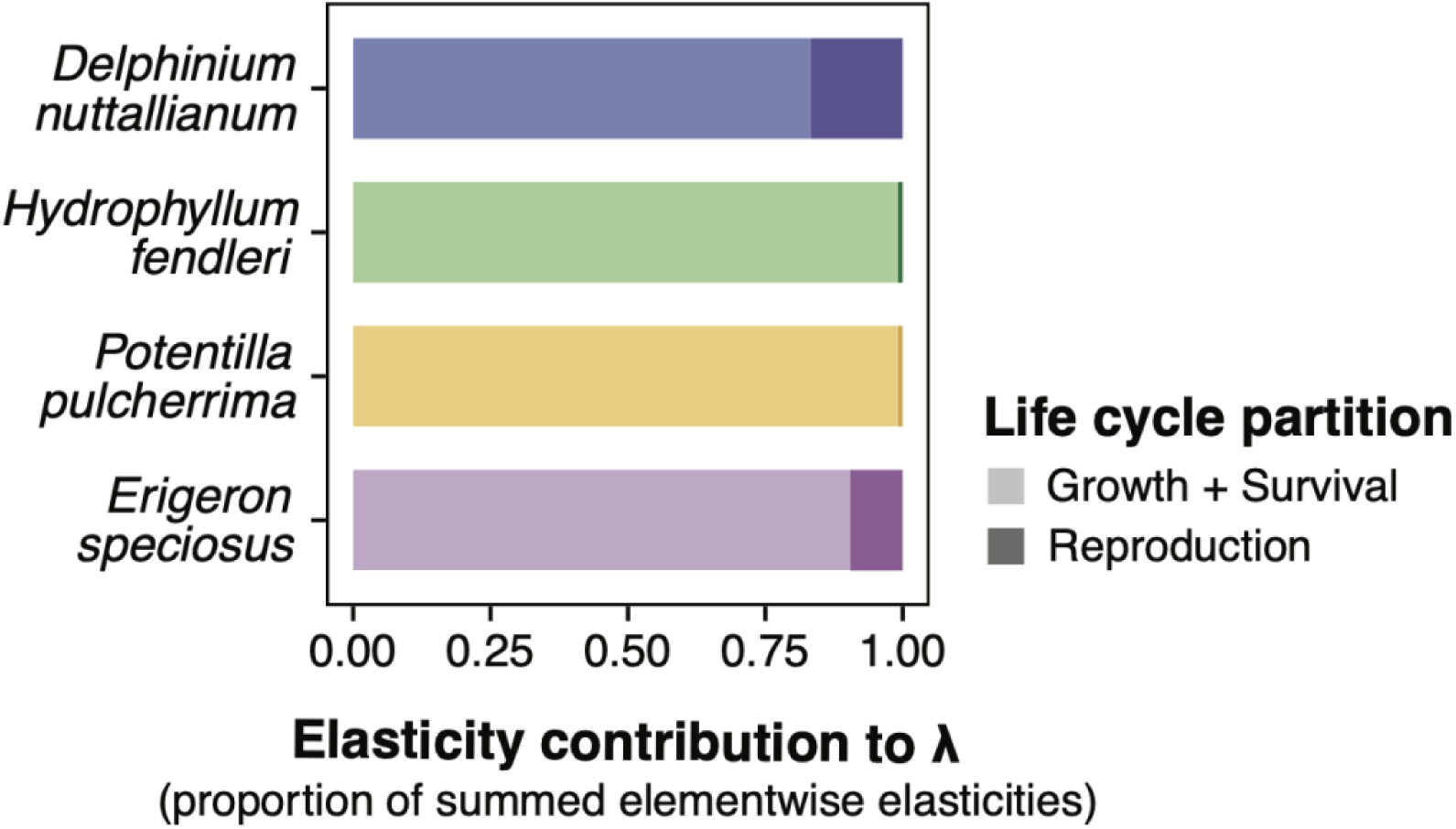
Median elasticity contributions to λ of the reproduction and persistence (growth and survival) partitions of the life cycle for each plant species (*Supporting Information, 2f*). Elasticity is the proportional change in λ in response to perturbations in different vital rates. (For the only species with a seed bank, *Hydrophyllum*, we included survival in the seed bank in the persistence partition and seed bank entry and emergence as reproduction.) Within each species, the summed elasticities from perturbations to growth and survival were higher than reproduction, indicating that changes to growth and survival most affect λ.

## Discussion

Our multi-species, experimental demography study provides empirical evidence that iteroparous perennial plant populations, which make up most plant biodiversity on the planet, can be buffered against population decline in the face of partial loss of pollination services. More generally, our results validate the predictions of demographic life history theory that argue for a whole life-cycle approach to measuring fitness impacts. Decreased pollination reduced seed production but only caused population decline in one species. Increased seed production in response to increased pollination came at a cost to survival that caused a net *reduction* in λ relative to the control in three species. Our experiment additionally exemplifies how changes in reproduction can fail as proxies for population-level outcomes. We would have predicted the correct *direction* of change in λ in only one half of our altered pollination conditions if we had relied on plant-level seed production as a heuristic for population response, as has been done in many previous studies (*reviewed by* Crone 2001; Iler *et al*. 2021).

In only one case did a decrease in pollination lead to a reduction in plant population growth rates (*Delphinium*), and this reduction occurred largely via reduced seed production. Additionally, *Delphinium* control plants exhibited the largest proportional change in λ when recruitment was perturbed, showing that its more reproduction-dependent life history could make this species more vulnerable to perturbations to reproduction. *Delphinium* might also have a lower capacity for trade-offs compared to the other species; more of its biomass is unavailable for reallocation from reproduction because it allocates the highest proportion of its biomass to flowering (Table 1; *Supporting Information*, *2d*). *Delphinium*’s response illustrates how iteroparous perennial plants can be at risk of reduced population growth rates under scenarios of pollinator decline. However, this species was the exception.

Population growth rates of the other three species were buffered against the observed level of seed loss in the decreased pollination treatment (*Hydrophyllum, Potentilla*, and *Erigeron*), showing no net change in λ. This result suggests that iteroparous perennial plants may be more resistant to immediate population declines under decreases in pollination than is typically hypothesized. Two mechanisms were responsible for this buffering. First, demographic trade-offs allowed increases in survival and growth that offset negative effects of the decreased pollination treatment on λ. Second, the life histories of these three species makes λ most elastic to perturbations of survival, which in combination with trade-offs, explains why λ was largely buffered against perturbations to reproduction. We caution that this buffering capacity will have a limit beyond which additional seed loss cannot be compensated. The potentially widespread capacity of iteroparous perennials to buffer against population decline under partial declines in pollination services should motivate action to reverse pollinator declines before this buffering capacity is exceeded.

Under the increased pollination treatment, why did λ of three species *decrease*, despite increases in seed set? In *Delphinium, Hydrophyllum,* and *Erigeron*, the higher cost of reproduction under increased pollination reduced survival, a vital rate to which λ is highly elastic. Other studies show that increased pollination can lead to reduced probability of flowering, fewer flowers, and reduced size in future years—but these costs did not result in significant decreases in λ (Baer & Maron 2018; Ehrlén & Eriksson 1995). Plants have evolved to reproduce under stochastic pollinator availability (Burd 1995), and natural selection is expected to maximize reproduction within the limits imposed by demographic trade-offs (Calvo, 1993).

The chronic increases in pollination that we impose here might represent a novel environment to which plants have evolved few mechanisms to rein in the increased costs of reproduction. Our results show how within-individual costs of reproduction are responsible for net reductions in λ under increased pollination, but other demographic mechanisms can produce this same counterintuitive relationship. In a monocarpic perennial, λ decreases non-linearly with seed inputs, due to negative density-dependence at life stages following seedling establishment (Campbell *et al*. 2017; Price *et al*. 2008; Waser *et al*. 2010). Our results suggest that management interventions to increase seed production of rare or declining plants through hand pollination might impose costs that exceed the benefits in iteroparous perennials. More broadly, our results highlight the ability of non-reproductive responses to altered pollination to determine the fate of plant populations, something that is not obvious unless viewed from a demographic life-history perspective.

The only species to show any evidence of an increase in λ in response to increased pollination, *Erigeron speciosus*, also showed the largest increase in seed production in response to increased pollination. These results are consistent with *Erigeron* being the only species in our study that was self-incompatible (Ingold *et al*. 2024) and therefore the most likely to benefit from receipt of additional outcross pollen. For changes in seed production to affect λ directly, λ must be somewhat elastic to perturbations in fecundity (*e.g.,* Law *et al*. 2010), which is the case for *Erigeron*. Additionally, the *Erigeron* population was growing under ambient pollination: λ = 1.20, a 20% annual increase in population size. Growing populations are more responsive to changes in reproduction because exponential growth magnifies even small increases in recruitment (Law *et al*. 2010; Silvertown *et al*. 1993). Finally, another reason there was some evidence that λ increased in response to increased pollination is that costs of reproduction were too low to totally offset the gains from increased seed production, as has been found in other species (Calvo 1993; Parker 1997).

The absence of a consistent link between decreased pollination and declines in plant population growth rates in our study has implications for understanding the potential effects of pollinator declines on species coexistence and species interaction networks. For pollinator declines to affect coexistence outcomes or the stability of plant-pollinator networks, changes in pollination must first affect plant population growth rates. This simplifying assumption is often acknowledged but rarely tested (Bartomeus *et al*. 2021; Bascompte *et al*. 2003; Memmott *et al*. 2004; Rohr *et al*. 2014; Thébault & Fontaine 2010). Experimental evidence from annual plants shows that pollinator declines can strengthen competitive imbalances between plant species pairs, weakening species coexistence, and potentially eroding plant biodiversity (Johnson *et al*. 2022). The buffering capacity we show of perennial plants against changes in pollination—a mechanism not possible in annual life histories—suggests that pollinator impacts on coexistence in perennial plant communities should result in less dramatic changes in competitive outcomes. Additionally, our results suggest that plant-pollinator network models should explore a range of plant population-level outcomes in simulations of pollinator extinction (as well as changes to pollination more broadly). In particular, the stability of λ in response to decreased pollination in three of our four species suggests that networks are likely to be more robust to pollinator loss than previously shown. These implications represent promising areas for future research, as our study only represents four plant species.

Extreme pollinator declines and complete pollinator extinctions will eventually lead to plant population declines when the buffering capacity shown in our study is exceeded. Observational evidence suggests that iteroparous, perennial plant species are already declining in response to pollinator declines (Anderson *et al*. 2011; Biesmeijer *et al*. 2006). Consistent with this, insect-pollinated plants have declined in sites across northern Europe (Ehlers *et al*. 2021; Pan *et al*. 2024). Parallel declines in plants and pollinators does not necessarily mean that pollinator declines are driving plant declines; however, outcrossing species and those relying on specialized pollination show the most extreme declines, which is quite suggestive (Biesmeijer *et al*. 2006; Ehlers *et al*. 2021). Moving forward, we need more studies that link changes in pollination to plant population growth rates, including experiments that vary the intensity of reduced pollination to find the limits of the buffering capacity of λ. Pairing experiments with time series or historic vs. recent comparisons that show declines in both plant and pollinator populations would be especially powerful.

Our results *do not* imply that pollinator declines and pollination are unimportant when they do not immediately cause plant population decline. Loss of fruits and seeds from an ecosystem could affect frugivores and granivores, crop production, and seed dispersal (Llanos-Guerrero *et al*. 2024). Furthermore, many pollinators transfer outcross pollen that ultimately increases the genetic diversity of plant populations. Finally, populations of some plant species are more likely to be vulnerable to losses in pollination than others. Species most at risk likely include annuals, semelparous perennials, and some iteroparous perennials, particularly those that meet some or all of the following: (i) the magnitude of seed loss is large, (ii) λ is highly elastic to changes in seed production, (iii) demographic trade-offs either do not exist or are too weak to compensate for seed losses, (iv) recruitment is seed limited (*e.g.*, Anderson *et al*. 2011; Lundgren *et al*. 2015), and (v) clonal reproduction is absent (Lin *et al*. 2016; Pauw & Bond 2011). The magnitude of seed loss in response to pollinator declines will in turn depend on the magnitude of pollen limitation and the contribution of pollinators to seed production (Rodger *et al*. 2021). This is exemplified in our study by *Potentilla*, the only species capable of autogamous self-pollination, which experienced only a 15% decline in seed production in response to a 50% or greater decrease in flowers available for animal pollination.

There is still much to be learned about the population-level consequences of mutualistic interactions, which remain chronically understudied (Ehrlén *et al*. 2016; Morris *et al*. 2020). Forecasting the impact of pollinator declines on plant persistence will require incorporating observed magnitudes of change in pollination that arise from actual pollinator declines. Nonetheless, our study addresses an important gap in our understanding of the implications of pollinator declines for plant populations and improves our general understanding of how mutualistic interactions affect population-level processes.

## Acknowledgements

We are thankful to the many field assistants, students, and teachers who collected data for this project. Special thanks to J Jonas and M Jones for their dedication and fieldwork contributions. The Iler-CaraDonna Lab, A. Compagnoni, D. Simpson, J. Beck, J. Thomson, M. Price, N. Waser, and S. Wagenius provided insightful feedback on earlier drafts of this manuscript. Volunteers from the Chicago Botanic Garden assisted with seed counting: C. Rolfini, D. Jaeschke, D. Tuerk, J. Newfeld, M. Frey, S. Higgins, S. Tepper. I. Rodelius, J. Ogilvie, V. DeLira, and numerous Northwestern University undergraduate students also assisted with seed counting. S. Levin and J.B. Socolar aided in model construction. J. Reithel and the Rocky Mountain Biological Laboratory (RMBL) provided access to field sites. M. Dorf participated in the RMBL Art-Science Exchange and encouraged us to think more expansively about our work.

## Author contributions

All authors contributed to the conceptualization, methodology, and acquisition of funding. A.M.I. led data acquisition, data curation, project management, and wrote the initial draft. P.J.C. led data visualization in addition to supporting the data acquisition, and project management. W.K.P. constructed, parameterized, and analyzed the models in addition to supporting data curation. All authors contributed to manuscript revisions.

## Data accessibility

Data and EML metadata are archived at the Environmental Data Initiative (package id: edi.2125.1; doi:10.6073/pasta/f4647d74c795548bf4e5237e85e6d41c). The code and data artifacts to reproduce the vital rate models, IPMs, and LTRE analyses are archived at Zenodo (doi: 10.5281/zenodo.15428278; full access available at: https://zenodo.org/records/15428278?token=eyJhbGciOiJIUzUxMiIsImlhdCI6MTc1NzU0NjMyNiwiZXhwIjoxNzY3MTM5MTk5fQ.eyJpZCI6IjY3NmU3NjBlLWQ4MTItNGZlNy1hN2ZhLWJlYzUwMDUxMWQxMCIsImRhdGEiOnt9LCJyYW5kb20iOiJjMWE3M2U4Zjc1ZGMyOGM4OTFmY2Y5M2M3MzQyNTRiYSJ9.knrizI71ifwX5ucQHZ_KeKvLn7_xx2X6ukOBC_uJ6s4E70Ec8S2UUee7tE7qQEm_fRaK5DBPYVrjRWBnk4hAZQ).

