## Supplemental Information for "Demographic trade-offs decouple pollination services from plant population growth"

**This file includes:**

Supporting text
- section 1: Building intuition for sensitivity
- section 2: Supplemental Materials & Methods

Figures S1 to S17
Table S1
SI References

### Supporting Information Text

#### 1. Building intuition for the sensitivity of the per-capita population growth rate to vital rate perturbations

A population at its stable distribution grows at a per-capita rate, $\lambda$ (Liu *et al.* 2011). In addition to this common population dynamic interpretation, $\lambda$ is equal to the expected number of offspring each individual contributes to subsequent generations over the course of its lifetime (*i.e.,* average fitness). Population size will grow, shrink, or remain stable when the average individual's lifetime offspring production exceeds, falls short of, or exactly equals replacement rate, respectively. Understanding why the fitness of individuals may be sensitive to (or buffered against) perturbations of a demographic vital rate can thus explain the sensitivity of population dynamics.

Life history determines the sensitivity of $\lambda$ to perturbations of the underlying demographic vital rates. The per-capita population growth rate will be strongly determined by the annual rate of offspring production in species that concentrate reproduction into few bouts during their lifetime. This may be due to short lifespan (*e.g.*, annuals) or a reproductive strategy that concentrates all reproductive investment into a single bout at the end of life (*i.e.,* semelparity). In contrast, species with long-lived, iteroparous life histories (*i.e.,* perennials) have per-capita population growth rates that are much more sensitive to perturbations of the survival rate than of the annual offspring production rate.

Here we use a toy example to first illustrate the population-level phenomenon of the sensitivity of $\lambda$ to perturbations of an iteroparous perennial organism. We then re-cast these perturbations from the perspective of individual-level demographic performance to build intuition. We refer readers who are interested in building their intuition further to Franco & Silvertown (2004) for comparative analysis of plant life histories based on formal analysis of the elasticity of $\lambda$ to perturbations in the vital rates, which is a proportional change in $\lambda$ (De Kroon *et al.* 1986).

***A simple population model for an iteroparous perennial***

Consider the life cycle of an iteroparous perennial for which the demographic vital rates depend on size and individuals reach reproductive maturity upon entering the largest class,

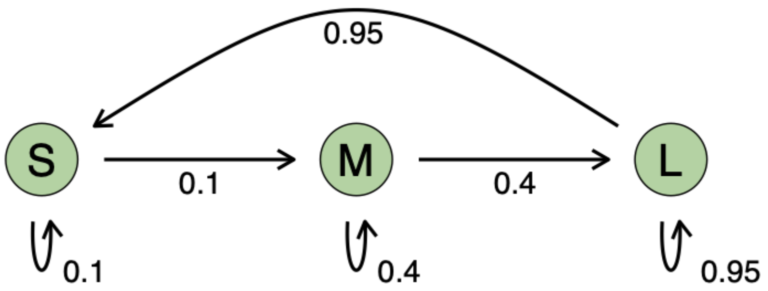


We can express this life cycle diagram equivalently as the matrix population model,

$$\begin{aligned}\mathbf{A} &= \mathbf{U}+\mathbf{F}\\ &= \begin{bmatrix}0.1 & 0 & 0\\ 0.1 & 0.4 & 0\\ 0 & 0.4 & 0.95\end{bmatrix}+\begin{bmatrix}0 & 0 & 0.95\\ 0 & 0 & 0\\ 0 & 0 & 0\end{bmatrix} \\ &= \begin{bmatrix}0.1 & 0 & 0.95\\ 0.1 & 0.4 & 0\\ 0 & 0.4 & 0.95\end{bmatrix}.\end{aligned} \qquad (1)$$

Individuals in the large size class have a 95% probability of persisting to the next year and produce, on average, 0.95 new offspring each year. For this population, λ = 1.017 corresponding to a modest 1.7% annual population increase.

***Sensitivity of λ to vital rate perturbations: offspring production & survival***

We can impose a perturbation that reduces the annual *rate of offspring production* by 1/3, by substituting the following for the original **F** matrix,

$$\mathbf{F}^* = \begin{bmatrix}0 & 0 & \mathbf{0.633}\\ 0 & 0 & 0\\ 0 & 0 & 0\end{bmatrix}, \qquad (2)$$

while leaving the **U** matrix unmanipulated. This means that every individual experiences the reduced offspring production rate for every reproductive bout of its lifetime. The resulting projection matrix is

$\mathbf{A}^* = \mathbf{U} + \mathbf{F}^*$. Despite the apparent severity of this perturbation, λ = 0.997. The population is declining towards extinction, albeit slowly at 0.3% per year.

Next, suppose that the perturbation instead reduced the *survival rate* of the largest size class, but with equal magnitude to the previous perturbation of the annual reproductive rate (33% reduction). Here, we can adjust the **U** matrix to,

$$\mathbf{U}^* = \begin{bmatrix} 0.1 & 0 & 0 \\ 0.1 & 0.4 & 0 \\ 0 & 0.4 & \mathbf{0.633} \end{bmatrix}, \qquad (3)$$

while leaving the **F** matrix unmanipulated, and re-project the resulting transition matrix. The resulting projection matrix is $\mathbf{A}^{**} = \mathbf{U}^* + \mathbf{F}$. Here, λ = 0.780 corresponding to a rapid decline towards extinction at 22% per year.

***Individual-level demographic responses to perturbations***

Why does the same perturbation to these two initially equal vital rates have such profoundly different consequences for population dynamics? Demographic performance during the pre-reproductive life stages is identical across all three matrix models, which means we can focus exclusively on the prospect for future life and reproductive bouts for an individual that reaches maturity.

***Individual-level distribution of the reproductive lifespan***

Upon reaching reproductive maturity, the future lifespan of individuals follows a geometric distribution (Fig. S1). The expected time to death is $\mathrm{E}(X) = 1/(1-s)$, where $s$ is the annual survival rate of reproductively mature individuals. For the unmanipulated model, **A** (Eq. 1), a newly maturing individual can expect to have 20 years of life and bouts of reproduction. The survival rate is unchanged in $\mathbf{A}^*$ (Eq. 2), meaning that the distribution of reproductive lifespan is unchanged. In contrast, the reduced survival rate in $\mathbf{A}^{**}$ (Eq. 3) causes the expected reproductive lifespan to fall to 2.73 years.

***Individual-level distribution of lifetime reproduction***

Taken one step further, we can simulate the lifetime reproduction of a cohort of individuals that have just reached reproductive maturity under each vital rate scenario by multiplying the reproductive lifespan by the annual rate of offspring production. Here too, the impact of reducing the annual survival rate overwhelms the impact of the reduced annual rate of offspring production (Fig. S2).

***Key takeaways***

For iteroparous life histories, dying early cuts all future offspring production to zero, which is a more severe perturbation than losing only a fraction of that future offspring production. Conversely, short lived and semelparous life histories do not afford individuals many (if any) opportunities for future offspring production. Therefore, in short lived and semelparous life histories, $\lambda$ will be sensitive to the annual offspring production rate, but less sensitive to survival.

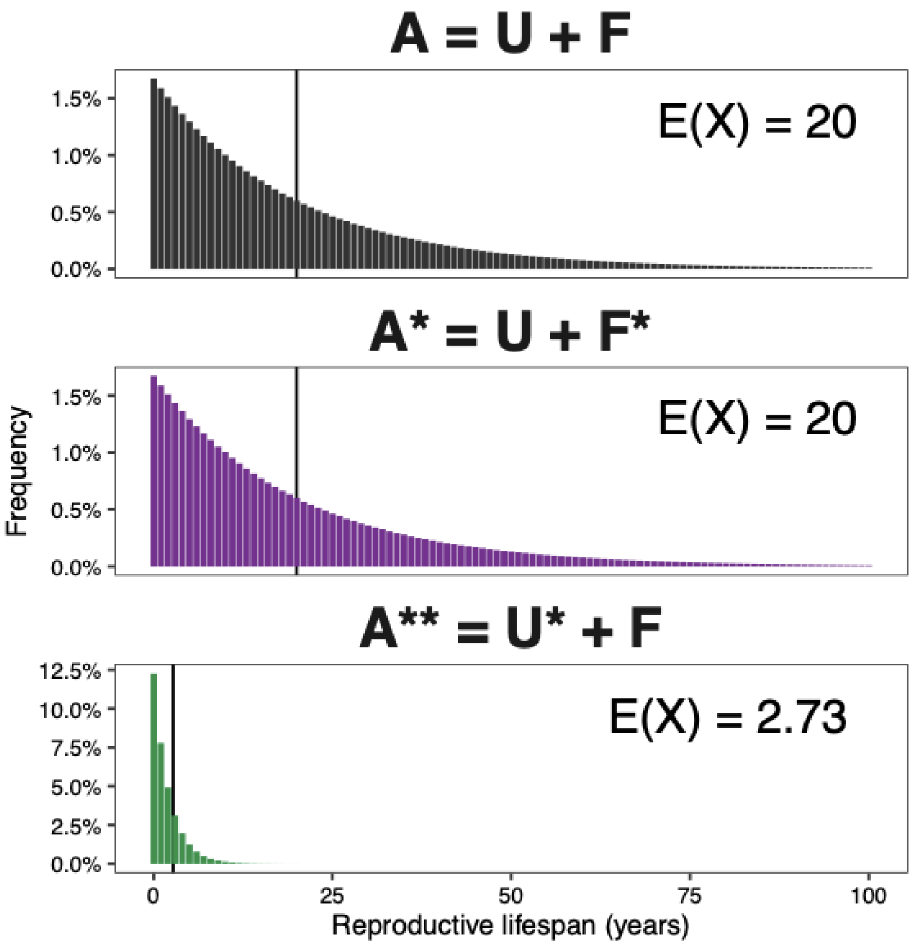


**Fig. S1.** Reducing survival by reduces reproductive lifespan much more than reducing offspring production by the same amount. Distribution of reproductive lifespan for the unperturbed matrix (top), a matrix in which offspring production is reduced by 1/3 (middle), and a matrix in which survival is reduced by 1/3 (bottom).

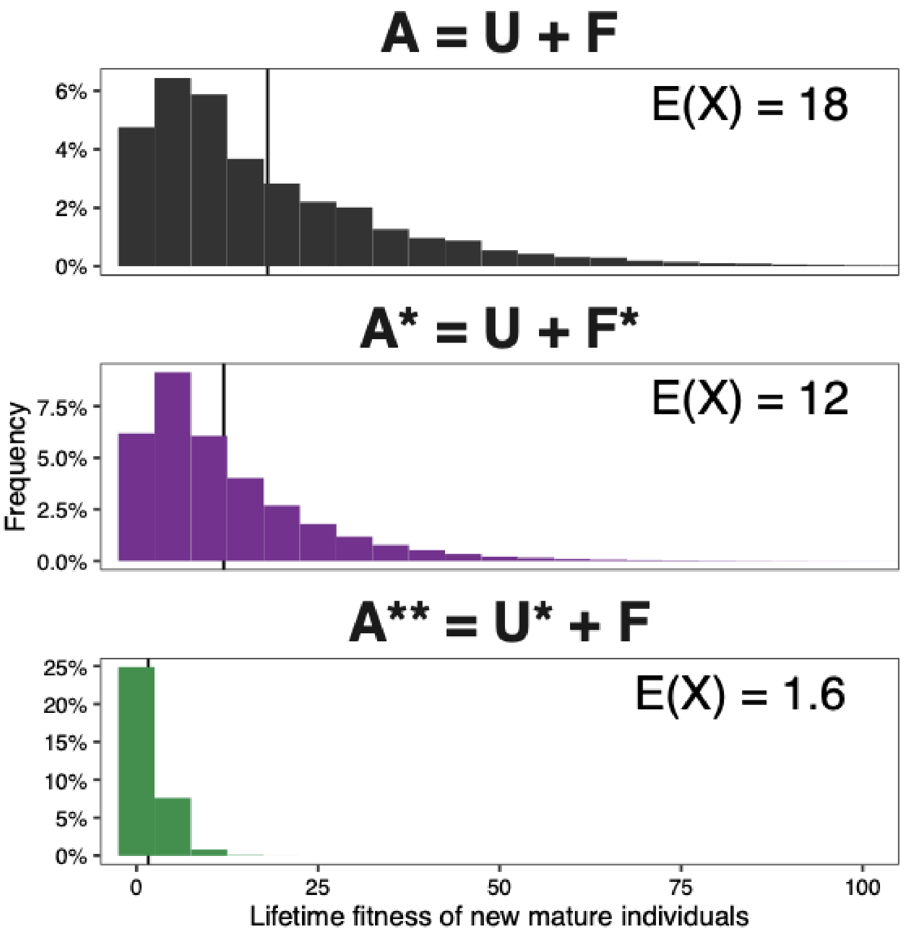


**Fig. S2.** Reducing survival reduces lifetime reproduction much more than reducing offspring production by the same amount. Distribution of lifetime reproduction for individuals upon reaching reproductive maturity for an unperturbed matrix (top), a matrix in which offspring production is reduced by 1/3 (middle), and a matrix in which survival is reduced by 1/3 (bottom). Each panel simulated a cohort of 1 million individuals upon reaching reproductive maturity.

### 2. Supplemental Materials and Methods

#### *2a. Study species extended*

Two of our study species, *D. nuttallianum* and *P. pulcherrima*, occur primarily in open meadows; *E. speciosus* occurs in both meadows and aspen understory, and our study population was in the latter habitat; *H. fendleri* occurs primarily in aspen understory and extends slightly into nearby open meadows; our study populations included both habitats. The community of native pollinators is largely intact and includes bumble bees, solitary bees, flies, hummingbirds, butterflies, and moths: there are no non-native honeybees (*Apis mellifera*), whose presence can affect foraging behavior of native bees. The four study species were selected because they are all animal-pollinated, herbaceous perennials that vary in phylogeny, phenology, and reproductive traits (Table 1, main text; Fig. S3). We expected plant mating system to be important for determining the degree to which altered pollination environments affected seed production because plant mating systems determine the degree to which a plant species depends on pollinators to set seed. The different flowering dates of the species allowed us to study them sequentially throughout the season.

*Delphinium nuttallianum* has a geographic range across western North America like all our study species, ranging from northern Arizona and New Mexico, USA in the south to British Columbia and Alberta, Canada in the north (USDA plants database). Populations of *D. nuttallianum* can be found from approximately 1,200 – 3,200 m a.s.l. (Waser & Price 1990). Plants have 1–14 palmately lobed leaves in our study population (mean 2 leaves) and do not reproduce vegetatively (Waser 1978). *Delphinium nuttallianum* plants produce one or occasionally two racemose inflorescences with 1–22 flowers in our study population (mean 4 flowers). The blueish-purple flowers are zygomorphic and protandrous and contain a nectar spur (Fig. S3A). Plants are self-compatible and therefore can produce seed with the receipt of self pollen, but seed production requires an animal pollinator to transfer pollen from one flower to another (Waser & Price 1990, 1991). Plants in our study population produce anywhere from 1 – 380 seeds, with an average of 33.8 seeds. Plants flower from late May through early July in our study area (Waser 1978). Flowers are primarily pollinated by hummingbirds (primarily the Broad-tailed hummingbird, *Selasphorus platycercus*) and bumble bee queens (Waser & Price 1981, 1990). This species does not have a seed bank (Waser & Price 1985).

The geographic range of *H. fendleri* spans from southern New Mexico in the USA northward to British Columbia, Canada (USDA plants database). Plants have 1 – 352 pinnately lobed leaves in our study population (mean 7 leaves). *Hydrophyllum fendleri* can spread vegetatively via short rhizomes; connections can be felt by gentle excavations of the soil surface (Iler, CaraDonna, & Petry, personal observation). Reproductive plants produce 1–121 raceme-like cymes (inflorescences) per plant (mean 3.0 cymes), each containing approximately 5–30 flowers (Fig. S3B). Plants are self-compatible, like *D. nuttallianum*, and therefore can produce seed with the receipt of self pollen but require an animal pollinator to transfer pollen from one flower to another (Beckmann Jr 1979). Plants flower from approximately June through July in our study area, and flowers are primarily visited by bumble bees (Iler & CaraDonna, unpublished data).

*Potentilla pulcherrima* has a geographic range spanning from southern Arizona and New Mexico in the USA through the southern provinces of west-central Canda (USDA plants database). In our study population, individuals produce 1–262 leaves that are palmately compound (mean 15 leaves), and individual plants produce flower stalks that each contain 1–177 yellow, actinomorphic flowers (mean 22 flowers; Fig. S3C). *Potentilla pulcherrima* plants can reproduce vegetatively (Burkle & Irwin 2010), and rhizomes can be gently excavated to identify individuals (Iler, CaraDonna, & Petry, personal observation). Plants are capable of autogamous self-pollination but produce more seeds if pollinated by insects and also produce more seeds with receipt of outcross vs. self pollen (Burkle & Irwin 2010). Plants typically flower from late June through early August in our study area. Flower visitors include syrphid flies, flies, solitary bees, and butterflies in our study area (CaraDonna *et al.* 2017).

*Erigeron speciosus* has a geographic range that extends across 11 states of the western USA and western Canada (CL Gucker & NL Shaw 2018) and an elevation range that spans 600–3,400 m a.s.l. Individual plants produce 1–171 stalks with multiple, elliptical leaves (mean 8 stalks, Fig. S3D). Each stalk typically contains a radiate inflorescence (or several inflorescences) with lavender ray flowers and yellow disc flowers (Fig. S3D), and the root system includes rhizomes and a caudex (CL Gucker & NL Shaw 2018). Plants are self-incompatible and therefore require transfer of outcross pollen from neighboring plants to set seed (Ingold *et al.* 2024). *Erigeron speciosus* typically flowers from mid-July through August in our study area and is visited by a wide variety of visitors, including bumble bees, several species of solitary bees, syrphid flies, bombyliid flies, and Lepidopterans (Burkle & Irwin 2009; CaraDonna & Waser 2020; Kearns 1992; Pohl *et al.* 2011).

***2b. Species-specific details of seedling emergence experiments***

*Delphinium nuttallianum*

For *D. nuttallianum*, we used plots of a constant size and varied seed sowing densities to create the seed density treatments. We established plots with a 12.5 cm radius on 15 July 2019. There were a small number of fruits (< 20 in entire area) that had opened on individuals growing within the buffer zone when we established the plots. All fruits were collected by 26 July, and seeds were sown on 2 August. We first tried to base seed densities on the number of seeds from the previous year. However, seed production was very low in 2018, presumably due to drought conditions (Faust & Iler 2022). To make sure we had enough variation in seed densities, we therefore based seed production on previous research; we assumed plants make three fruits per plant in a 'good year' (NM Waser & MV Price, personal communication). To set the lower bound of the density treatments, we multiplied mean seeds per fruit from the removal treatment by 3 fruits. To set the upper bound, we multiplied mean seeds per fruit from the supplemental treatment by 3 fruits, then by 10 (1160 seeds per $m^2$). The final seed densities were: 0 (control), 5, 13, 22, 36, and 58 seeds per plot, replicated three times for a total of 18 plots, and 248 seedlings emerged in 2020, for a mean germination rate of 61.7%. An average of 8.33 seedlings emerged in control plots, showing that there was some natural seed rain into the plots. We used a generalized linear model (GLM) with a negative binomial error distribution and sowing density as a continuous predictor to analyze the number of seedlings (using package MASS, *(WN Venables & BD Ripley 2002)*.

*Hydrophyllum fendleri*

For the following three species (*H. fendleri, P. pulcherrima,* and *E. speciosus*)*,* we varied the size of the plots and held seed inputs constant (100 seeds per plot) to create the seed density treatments. We used a different design for the remaining three species because we observed low germination rates in the demography study and wanted to maximize the chance of detecting seedling emergence events. For all species, we sowed only seeds that appeared to be viable (*i.e*., plump and full) that we collected from plants within the study populations, but outside demography plots.

The following plot radii were used for *H. fendleri*: 11 cm, 15 cm, 20 cm, 24 cm (control), 26 cm, and 48 cm. Plots were established and seeds sown on 5–7 August 2019. No seeds germinated in 2020 (from seeds sown in 2019), so we repeated this experiment in 2020. In 2020, on 13 August, we lightly buried the seeds just below the soil surface when we sowed them. Sixteen seedlings emerged in 2021 across all plots, for a mean germination rate of 0.88%. Following the GLM for *D. nuttallianum*, the number of seedlings were analyzed using a GLM with a negative binomial error distribution and sowing density as a continuous predictor.

*Potentilla pulcherrima* & *Erigeron speciosus*

The following plot radii were used for both *P. pulcherrima* and *E. speciosus*: 9 cm, 11 cm, 14 cm, 16 cm (control), 16 cm, 20 cm, and 24 cm. Plots were established and seeds dispersed on 2 October 2019 for *P. pulcherrima* and 7 October 2019 for *E. speciosus*. Only one replicate was established in 2019 as a pilot because flowers were not removed within a 1 m radius of each plot; no seeds germinated in 2020. We repeated this experiment in 2020, increased the replicates to three per treatment, and we lightly buried the seeds just below the soil surface. For *P. pulcherrima*, the remaining plots were established and flowers removed on 30–31 July 2020, and seeds were sown on 18 August 2020. For *E. speciosus*, the remaining plots were established and flowers removed on 11–12 August 2020, and seeds were sown on 1 September 2020.

***2c. Seed bank experiment***

The existence of a soil seed bank poses challenges for population models because seed banks partially decouple plant abundances between years (Crawley 1990). *Delphinium nuttallianum* does not have a seed bank in our study area (Waser & Price 1985), but the seed bank status of our other three study species was unknown. We therefore buried known quantities of seed in nylon mesh bags to determine the proportion of seeds that germinate one-, two-, and three-years after burying in the soil (2019–2021). We collected seeds from plants adjacent to the experimental demography plots and within the natural bounds of each study population. We used 30 nylon mesh bags per species (10 per year), and seed densities in the bags were based on seed production rates and flowering plant density in control plants in 2018 (5 seeds per bag for *H. fendleri*, 10 seeds per bag for *P. pulcherrima*, and 13 seeds per bag for *E. speciosus*). We filled each bag with soil from the site, added seeds in August 2018, and buried the bags adjacent to the demography plots, with approximately 1 cm of dirt covering each bag. In only one species did we find evidence of a seed bank, *H. fendleri*, with a 19.3% germination rate in year one, a 12.4% germination rate in year two, and an 11.3% germination rate in year three. Details of how we incorporated seed bank dynamics into the IPM are below. We observed seedlings of *P. pulcherrima* and *E. speciosus* in the bags only in year one, at a rate of 4% and 4.6%, respectively. We therefore did not need incorporate a seed bank into IPMs for these species.

***2d. Proportion of biomass allocated to reproductive effort***

The proportion of biomass allocated to reproductive effort provides insight into resources available for reallocation to vital rates to which λ is most sensitive (*i.e.,* survival and growth), following changes in seed production. We excavated individuals of our four study species from areas adjacent to our experimental demography plots and seedling emergence plots. Plots of various size were established to allow sampling of 20 flowering and 20 non-flowering individuals of each species; only the flowering individuals are used here. The *D. nuttallianum* plot was 13 m x 17 m, the *P. pulcherrima* plot was 5 m x 17 m, and the two *E. speciosus* plots were 5 m x 9 m and 5 m x 10 m. A random number generator was used to generate 20 coordinates within each plot. The closest, flowering plant to those coordinates was excavated from the ground, using standard gardening trowels and shovels. Due to space constraints, *H.*

*fendleri* plants were extracted from random coordinates around the perimeter of each of the 16 demography plots.

Individuals were removed from the ground either during peak fruiting (*D. nuttallianum* and *H. fendleri*) or peak flowering (*P. pulcherrima* and *E. speciosus*). The plants were dried in a drying oven for at least 48 hours and weighed for total biomass, above-ground biomass, below-ground biomass, and reproductive biomass (flowering stalks and flowers, if present at the time of collection) on a microbalance to the nearest 0.001g.

We calculated the percent of reproductive biomass out of total biomass by dividing reproductive biomass by total biomass and multiplying by 100 (table S1). Because *D. nuttallianum* was the only species that was 100% in the fruiting phase at the time of excavation, we calculated its total reproductive biomass as only the flowering stalk; this is an underestimation of biomass allocated to reproductive effort, but we wanted to ensure that fruit weights were not biasing our results. Even if reproductive biomass of *D. nuttallianum* is only represented by the flowering stalk with no fruits or flowers present, it still allocates more than twice the percentage of its biomass to reproductive effort than the other species (17.4% vs. values in Table 1 of main text).

#### *2e. Integral Projection Models and LTREs*

We built a size-structured, density-independent integral projection model (IPM) to infer each plant species' population dynamics under each pollination treatment. The abundance of individuals across the size range in the next time step, $\mathrm{n(z', t+1)}$, is calculated from the recursion equation,

$$n(z', t+1) = \int_L^U [P(z', z) + F(z', z)]n(z, t)dz. \qquad (1)$$

Here, the *kernel* describes how the future population arises from the current abundance-size distribution, $\mathrm{n(z', t+1)}$. The kernel is composed of two subkernels: $\mathrm{P(z', z)}$ and $\mathrm{F(z', z)}$, which describe how current plants lead to future plants by the persistence of established plants and/or the production of new offspring, respectively. $P$ and $F$ are expressed using functional notation with $z$ used to indicate a plant's current size and $z'$ to indicate its size in the next time step. This allows persisting plants to grow (or shrink) and for reproductive plants that are typically large to produce recruits that are typically small. Moreover, *any* functional form can be used to map current $z$-sized plants to future $z'$-sized plants, which allowed us to tailor the model for each plant species' size-specific demographic processes.

Each of the subkernels may be further broken down into components of the persistence or offspring production process. These components are the *demographic vital rates*. We modeled persistence as the product of size-specific survival, $s(z)$, and growth, $g(z', z)$:

$$P(z', z) = s(z)g(z', z). \qquad (2)$$

This subkernel impacts both the abundance and size distribution of the population at the next time step. Plants that die necessarily lower the total abundance and may also affect the relative frequency of sizes if the survival rate depends on size. The growth component does not affect the total population abundance but does affect the relative frequency of sizes by reallocating the individuals that do survive and grow (or shrink).

The offspring production subkernel can be similarly decomposed into the product of reproductive vital rates,

$$F(z', z) = f(z)b(z)r_r r_d(z'), \qquad (3)$$

where $f(z)$ is the size-specific probability of flowering, $b(z)$ is the expected number of seeds produced by a $z$-sized individual, $r_r$ is a size-independent probability of recruitment, and $r_d(z')$ is the recruit size distribution.

We tailored the component functions of each kernel to fit the life cycle of each species. The species-specific equations for each function and any deviations in kernel structure required to match the species' life history are described below (see section: *Species-specific demographic vital rate and Integral Projection Model methods*).

*Estimation of the demographic vital rate parameters of the IPMs*

We fit a single probabilistic hierarchical model to data from (i) the demographic census, (ii) the seed germination experiment, and (iii) the seedbank persistence experiment to characterize the size-and treatment-specific annual vital rates for each plant species. Fitting a separate model for each species allowed us to tailor different functional forms of the underlying vital rate functions to the biology of each species.

We fit the vital rate model for each species using Bayesian methods because this allowed us (i) to tailor the multiple response variable distributions to fit the data-generating process (vs. assuming multivariate normal), (ii) to estimate correlated intercept changes among response variables owing to plot and year effects so we could better estimate the demographic impact of resource allocation constraints, (iii) to avoid overfitting through the use of regularizing priors on treatment and size-by-treatment effects, and (iv) to account for how uncertainty in our parameter estimates affects our projections of population dynamics.

The generic vital rate model is given as,

$$
\begin{aligned}
&\text{Survival probability} && \begin{cases} Y_i^s & \sim \text{Bernoulli}(p_i^s) \\ \text{logit}(p_i^s) & = \beta_0^s + \beta_{trt}^s + \beta_z^s z_i + \beta_{z:trt}^s z_i + \alpha_{plot[j]}^s + \alpha_{year[k]}^s \end{cases} \\
&\text{Growth} && \begin{cases} [Y_i^g | Y_i^s = 1] & \sim \text{Normal}(\mu_i^g, \sigma_i^g) \\ \mu_i^g & = \beta_0^g + \beta_{trt}^g + \beta_z^g z_i + \beta_{z:trt}^g z_i + \alpha_{plot[j]}^g + \alpha_{year[k]}^g \\ log(\sigma_i^g) & = \gamma_0^g + \gamma_{trt}^g \end{cases} \\
&\text{Flowering probability} && \begin{cases} [Y_i^f | Y_i^s = 1] & \sim \text{Bernoulli}(p_i^f) \\ \text{logit}(p_i^f) & = \beta_0^f + \beta_{trt}^f + \beta_z^f z_i + \beta_{z:trt}^f z_i + \alpha_{plot[j]}^f + \alpha_{year[k]}^f \end{cases} \\
&\text{Seed production} && \begin{cases} [Y_i^b | Y_i^f = 1] & \sim \text{GammaPoisson}(\mu_i^b, \theta_i^b) \\ \log(\mu_i^b) & = \beta_0^b + \beta_{trt}^b + \beta_z^b z_i + \beta_{z:trt}^b z_i + \alpha_{plot[j]}^b + \alpha_{year[k]}^b \\ \log(\theta_i^b) & \sim \gamma_0^b + \gamma_{trt}^b + \tau_{year[k]}^b \end{cases} \\
&\text{Germination probability} && \begin{cases} [Y_i^r | \text{recruit} = 1] & \sim \text{Binomial}(n_i, p^r) \\ \text{logit}(p^r) & = \beta_0^r \end{cases} \\
&\text{Recruit size distribution} && \begin{cases} [Y_i^d | \text{recruit} = 1] & \sim \text{Normal}(\mu^d, \sigma^d) \\ \mu^d & = \beta_0^d. \end{cases}
\end{aligned}
\tag{4}
$$

We allowed the group-level intercepts for plot and observation year to be correlated across the vital rate functions as,

Correlated varying effects

$$
\begin{cases}
\begin{bmatrix} \alpha^{s}_{plot[j]} \\ \alpha^{g}_{plot[j]} \\ \alpha^{f}_{plot[j]} \\ \alpha^{b}_{plot[j]} \\ \vdots \end{bmatrix} & \sim \text{MVNormal}\left( \begin{bmatrix} 0 \\ 0 \\ 0 \\ 0 \\ \vdots \end{bmatrix}, \Sigma_{plot} \right) \\
\Sigma_{plot} & = \mathbf{S}_{plot} \mathbf{R}_{plot} \mathbf{S}_{plot} \\
\mathbf{R}_{plot} & \sim \text{Student}(3, 0, \sigma_{plot}) \\
\mathbf{S}_{plot} & \sim \text{LKJ}(\eta_{plot}) \\
\begin{bmatrix} \alpha^{s}_{year[k]} \\ \alpha^{g}_{year[k]} \\ \alpha^{f}_{year[k]} \\ \alpha^{b}_{year[k]} \\ \tau^{b}_{year[k]} \\ \vdots \end{bmatrix} & \sim \text{MVNormal}\left( \begin{bmatrix} 0 \\ 0 \\ 0 \\ 0 \\ \vdots \end{bmatrix}, \Sigma_{year} \right) \\
\Sigma_{year} & = \mathbf{S}_{year} \mathbf{R}_{year} \mathbf{S}_{year} \\
\mathbf{R}_{year} & \sim \text{Student}(3, 0, \sigma_{year}) \\
\mathbf{S}_{year} & \sim \text{LKJ}(\eta_{year}),
\end{cases}
\tag{5}
$$

where ⁝ indicates additional plot- and year-level varying effects used for species-specific modifications of the generic vital rate model (see *Species-specific demographic vital rate and Integral Projection Model methods*). The subscripts indicate the replicate, and all letter superscripts indicate the vital rate (*i.e.,* they are not exponents). We tailored this generic vital rate model to fit the life cycle of each species. The modifications to the vital rate model for each species are detailed in the section below: '*Species-specific demographic vital rate and Integral Projection Model methods'*.

For each species' vital rate model, we chose weakly informative prior distributions for all model parameters (Gelman *et al.* 2008; Lemoine 2019). We aimed to constrain the model priors to ranges that were broadly compatible with biological feasibility, and we validated the suitability of these priors through prior predictive checks for each response variable (*i.e.*, drawing samples from the model without supplying it any data). We conducted a power-scaling sensitivity analysis on the univariate fit for each vital rate response variable using the priorsense package in R (v.1.1.0, Kallioinen *et al.* 2024). We broadened the dispersion of priors when prior-data conflicts for which we did not have strong, independent belief in our original choices. The large sample sizes of our census datasets resulted in a posterior distribution that was strongly dominated by the data with comparatively weak influence of the prior distributions.

We fit our models using Stan (v.2.36) via the R package brms (v.2.22.0, Bürkner 2017, 2018; Stan Development Team 2025). We used the No-U-Turn sampler for Hamiltonian Monte Carlo with 2000 warmup iterations and 2000 iterations used for inference on each of four chains. We assessed the suitability of our fits for inference by ensuring that there were no divergent transitions and that all chains converged to the same posterior distribution as indicated by a maximum of the rank normalized split-$\hat{R}$ of less than 1.05 for all parameters (Vehtari *et al.* 2021).

*Projecting the population dynamics of the IPMs*

We constructed IPM kernels using the R package ipmr (v.0.0.7, Levin *et al.* 2021). The duration of our demographic censuses (4–5 years, depending on the species) limited our analyses to deterministic population projections using a kernel that marginalizes over temporal (year) and spatial (plot) varying effects (Doak *et al.* 2005). We determined the lower and upper size domains of the model by extending each bound *ca.* 15% beyond the observed size range, limiting this to the upper bound when the lower bound was naturally bounded (*e.g.*, number of leaves cannot be below one). We discretized the IPM kernel by evaluating the integral using the midpoint rule at 400 equally spaced size bins. To avoid eviction of individuals from the model because their future size falls outside the bounded size range, we used truncated distributions (Williams *et al.* 2012).

Once the IPM kernels were constructed, we determined the equilibrium per-capita population growth rate, λ, by projection of at least 100 timesteps or until convergence (indicated by a $< 10^{-6}$ change in the value of $\lambda_t$ compared to $\lambda_{t-1}$).

We used the R package Rage (v.1.8.0, Jones *et al.* 2022) to calculate the generation time for the Control treatment as the time required for the population to increase by a factor of $R_0$, the net reproductive rate (Caswell 2001). We used age-from-stage methods (Caswell 2001) in the R package

Rage (v.1.8.0, Jones *et al.* 2022) to calculate the longevity of plants in the Control treatment as the age at which the survivorship, $l_x$, of a synthetic cohort dropped below 1%.

*Life Table Response Experiment*

We conducted a one-way fixed Life Table Response Experiment (LTRE; Caswell 2001*)* to determine how the change in per-capita population growth rate, Δλ, arose from the differences in the underlying vital rate parameters between treatment and control for each plant species. As a first order approximation, this decomposition may be expressed as,

$$\Delta\lambda = \lambda_{trt} - \lambda_{ctrl} \qquad (6)$$

$$= \sum_{i}^{k} s_i \Delta\theta_i,$$

where $s_i$ is the sensitivity of λ to local perturbation of the vital rate parameter $\theta_i$ and $\Delta\theta_i$ is the change in the underlying parameter in the pollination treatment relative to the control, $\theta_{trt,i} - \theta_{ctrl,i}$. We measured the sensitivity of λ to each vital rate $\theta_i$ using perturbations of an IPM kernel built using the parameter values at the midpoint between the treatment and control, $\theta_{mid,i} = (\theta_{trt,i} + \theta_{ctrl,i})/2$ [(Ellner *et al.* 2016), pp. 103-105]. We approximated the sensitivities of each vital rate parameter, $i$, using centered differences as,

$$s_i = \frac{\delta\lambda_{mid}}{\delta\theta_{mid,i}}$$

$$\approx \frac{\lambda_{mid}(\theta_i + \Delta) - \lambda_{mid}(\theta_i - \Delta)}{2\Delta}, \qquad (7)$$

where Δ is an arbitrarily small perturbation. We chose Δ using a common heuristic: the cube root of the machine precision on the computer used for our analysis $\approx 6.1 \times 10^{-6}$.

We verified that our first order approximation of the sensitivities was sufficiently accurate by ensuring that $\Sigma_{i=1}^{k} s_i \Delta\theta_i$ terms did in fact approximate the total Δλ. The median absolute biases were

<0.002 for all species-treatment combinations and were most often an order of magnitude smaller. Because the bias and error were very small, we did not calculate second or higher order derivatives.

The additive nature of these terms allows us to sum the LTRE contributions of multiple vital rate parameters to capture the net contribution of a biologically meaningful set. For example, summing the terms for the survival function slope and intercept to determine the overall contribution of treatment effects on the survival function to their net effect on λ.

*Propagating uncertainty*

Fitting the vital rate model for each species in a Bayesian framework provided a natural means of propagating uncertainty in the low-level vital rate parameter estimates to the resulting per-capita population growth rate, λ, and LTRE contributions of the vital rate parameters (Elderd & Miller 2016).

We calculated the posterior distribution of λ for each treatment by parameterizing the IPM with 1000 joint posterior samples from the vital rate model and projecting the population dynamics to equilibrium. Importantly, each parameter set was sampled from the *joint* posterior distribution of the vital rates, which embeds the covariances between parameter values. Samples with, for example, a low value for the survival intercept in the control treatment may tend to have lower (or higher) values for the size slope for the pollen supplementation treatment in the growth function. Because of this, the difference between distribution of $\lambda_{\mathrm{trt}}$ and the distribution of $\lambda_{\mathrm{ctrl}}$ will not necessarily equal the desired distribution of $\lambda_{\mathrm{trt}} - \lambda_{\mathrm{ctrl}}$.

We applied this same uncertainty propagation approach to the LTRE analysis. Briefly, this involved calculating (i) the parameter differences between treatment and control and (ii) the mean of the treatment and control parameters for 1000 joint posterior samples from the vital rate model. Then, we parameterized the IPM with each mean parameter set and calculated the sensitivity of $\lambda_{\mathrm{mid}}$ to small perturbations of each vital rate parameter. Finally, we multiplied these sensitivities to their corresponding treatment–control parameter differences to determine the posterior distribution of LTRE contributions.

*Species-specific demographic vital rate and Integral Projection Model methods*

***Delphinium nuttallianum:*** The germination plots experienced seed rain from adjacent plants. We modified the germination probability component of the generic vital rate model to treat observed seedlings as a mixture of germinants from the experimentally-sown and naturally-dispersed seeds,

$$\begin{aligned} Y_i^r &\sim \text{Poisson}(\lambda_i^r) \\ \lambda_i^r &= \Lambda^r A_i n_i \theta^r, \end{aligned} \qquad (8)$$

where $\Lambda^r$ is the latent density natural seed rain, $A_i$ is the area of the seed plot, $n_i$ is the number of seeds experimentally sown into the plot, and $\theta^r$ is the rate of seed germination. We assume that both sources of seeds have the same germination probability.

*IPM kernel definitions:*

$$\boldsymbol{P}(\boldsymbol{z}', \boldsymbol{z})$$

$$\begin{aligned} s(z) &= \text{logit}^{-1}(\beta_0^s + \beta_{trt}^s + \beta_z^s z_i + \beta_{z:trt}^s z_i) \\ g(z', z) &= \text{TruncNormal}_L^U(\mu^g, \sigma^g) \\ \mu^g &= \beta_0^g + \beta_{trt}^g + \beta_z^g z + \beta_{z:trt}^g z \\ \sigma^g &= \gamma_0^g + \gamma_{trt}^g \end{aligned} \qquad (9)$$

$$\boldsymbol{F}(\boldsymbol{z}', \boldsymbol{z})$$

$$\begin{aligned} f(z) &= \text{logit}^{-1}\left(\beta_0^f + \beta_{trt}^f + \beta_z^f z + \beta_{z:trt}^f z\right) \\ b(z) &= e^{(\beta_0^b + \beta_{trt}^b + \beta_z^b z + \beta_{z:trt}^b z)} \\ r_r &= \theta^r \\ r_d &= \text{TruncNormal}_L^U(\mu^d, \sigma^d) \end{aligned} \qquad (10)$$

*Integration rule:* midpoint

L = -3.513

U = 15.199

Number of meshpoints: 400

***Hydrophyllum fendleri:*** We used the number of leaves as the size metric for *H. fendleri*, meaning that an individual's size was constrained to positive integers ($z \in \mathbb{Z}^{+}$). We accommodated this difference by (i) log-transforming $z$ when it was a predictor in a vital rate equation and (ii) substituting truncated count distributions for the growth and recruit size distribution vital rate functions (*i.e.*, $z = 0$ leaves is not possible). We modified the generic vital rate model by substituting the following for these vital rate functions,

$$\text{Growth} \quad \begin{cases} \left[Y_i^g | Y_i^s = 1\right] & \sim \text{TruncNegBinomial}^{+}\left(\mu_i^g, \theta_i^g\right) \\ \mu_i^g & = \beta_0^g + \beta_{trt}^g + \beta_z^g z_i + \beta_{z:trt}^g z_i + \alpha_{plot[j]}^g + \alpha_{year[k]}^g \\ log\left(\theta_i^g\right) & = \gamma_0^g + \gamma_{trt}^g + \tau_{year[k]}^g \end{cases} \tag{11}$$

$$\text{Recruit size distribution} \quad \begin{cases} \left[Y_i^d | \text{recruit} = 1\right] & \sim \text{TruncPoisson}^{+}(\lambda^d) \\ \lambda^d & = \beta_0^d. \end{cases}$$

The $\tau$ varying effects for plot and year on the overdispersion parameter were included in the correlated varying effects (Eq. 5).

Data from the buried seed bags indicated that this species has a perennial soil seed bank. We included a single discrete stage to track the dynamics of the seed bank. We assume that all seeds that did not germinate the following spring entered the seedbank. The vital rate function for seed bank survival was given as,

$$\text{Seed bank survival} \quad \begin{cases} \left[X_i^q \leq Y_i^q \leq Z_i^q\right] & \sim \text{Binomial}\left(n_i^q, p_i^q\right) \\ \text{logit}\left(p_i^q\right) & = \beta_0^q + \beta_t^q t_i + \alpha_{block[j]}^q + \alpha_{seedplot[l]}^q. \end{cases} \tag{12}$$

We used interval censoring on the response variable to account for observation uncertainty for several bags in which germinated seeds could not be confidently assigned to a germination year. Because we ran only one seed bag experiment for multiple years, we were unable to distinguish the varying effect of year. Neither of the $\alpha^q$ varying effects were included in the correlated varying effects structure because the seed bag experiment could not be conducted directly in the census plots without disturbing the soil and potentially altering the vital rates of tagged plants. We assumed that the slope for time measured the rate of emergence from the seed bank.

*IPM kernel definitions:*

Our choice of leaf number as the size metric for *H. fendleri* meant that the vital rate functions could only be meaningfully evaluated at discrete size values. Although we could still parameterize the demographic model using estimates from a regression model as we did for the other species, the resulting demographic model is formally a matrix population model (MPM) because an IPM *sensu stricto* includes at least one continuous state variable (Moutouama *et al.* 2025). In practice, the integrals for these models are calculated numerically from a discretized kernel. We adopted a broader definition for IPMs to include the model constructed for *H. fendleri*, emphasizing the common features our model shares with IPMs *sensu stricto*: (i) the demographic model is represented computationally as a large-dimension square matrix, the elements of which contain transition rates between values of the state variable, and (ii) the transition rates are estimated from regression analyses rather than direct observations of a given state-to-state transition. The tools for constructing IPMs in the package *ipmr* (Levin *et al.* 2021) are agnostic to the biological meaning of the state variable. By carefully choosing the integration limits and mesh points, we were able to produce a discretized "kernel" for the *H. fendleri* demographic model that only included biologically plausible integer values of the state variable at the midpoint.

The addition of a seed bank for this species requires additional kernels that track entry, persistence, and exit from the discrete stage. The projection kernel therefore becomes,

$$\begin{aligned} K(z',z) &= P(z',z) + F(z',z) + D(q,z) + P_{bank}(q) + E(z',q) \\ D(q,z) &= f(z)b(z)(1-r_r) \\ P_{bank}(q) &= s_{bank} \\ E(z',q) &= r_r r_d(z'), \end{aligned} \tag{13}$$

where $q$ is the discrete state variable for the soil seed bank.

$$\boldsymbol{P(z',z)}$$

$$\begin{aligned} s(z) &= \text{logit}^{-1}(\beta_0^s + \beta_{trt}^s + \beta_z^s z_i + \beta_{z:trt}^s z_i) \\ g(z',z) &= \text{TruncNormal}_L^U(\mu^g, \sigma^g) \\ \mu^g &= \beta_0^g + \beta_{trt}^g + \beta_z^g z + \beta_{z:trt}^g z \\ \sigma^g &= \gamma_0^g + \gamma_{trt}^g \end{aligned} \quad (14)$$

$$\boldsymbol{F(z',z)}$$

$$\begin{aligned} f(z) &= \text{logit}^{-1}\left(\beta_0^f + \beta_{trt}^f + \beta_z^f z + \beta_{z:trt}^f z\right) \\ b(z) &= e^{(\beta_0^b + \beta_{trt}^b + \beta_z^b z + \beta_{z:trt}^b z)} \\ r_r &= \text{logit}^{-1}(\beta_0^r) \\ r_d &= \text{TruncPoisson}^+(\lambda^d) \end{aligned} \quad (15)$$

$$\boldsymbol{D(q,z)}$$

$$\begin{aligned} f(z) &= \text{logit}^{-1}\left(\beta_0^f + \beta_{trt}^f + \beta_z^f z + \beta_{z:trt}^f z\right) \\ b(z) &= e^{(\beta_0^b + \beta_{trt}^b + \beta_z^b z + \beta_{z:trt}^b z)} \\ r_r &= \text{logit}^{-1}(\beta_0^r) \end{aligned} \quad (16)$$

$$\boldsymbol{P_{bank}(q)}$$

$$s_{bank} = \text{logit}^{-1}\left(\beta_t^q\right) \quad (17)$$

$$\boldsymbol{E(z',q)}$$

$$\begin{aligned} r_r &= \text{logit}^{-1}(\beta_0^r) \\ r_d &= \text{TruncPoisson}^+(\lambda^d) \end{aligned} \quad (18)$$

*Integration rule:* midpoint

L = 0.5

U = 400.5

Number of meshpoints: 400 (+ 1 discrete seed bank stage)

***Potentilla pulcherrima****:* The generic vital rate model was used for *P. pulcherrima* (Eq. 4).

*IPM kernel definitions:*

$$\boldsymbol{P(z',z)}$$

$$\begin{aligned} s(z) &= \text{logit}^{-1}(\beta_0^s + \beta_{trt}^s + \beta_z^s z_i + \beta_{z:trt}^s z_i) \\ g(z',z) &= \text{TruncNormal}_L^U(\mu^g, \sigma^g) \\ \mu^g &= \beta_0^g + \beta_{trt}^g + \beta_z^g z + \beta_{z:trt}^g z \\ \sigma^g &= \gamma_0^g + \gamma_{trt}^g \end{aligned} \quad (19)$$

$$\boldsymbol{F(z',z)}$$

$$\begin{aligned} f(z) &= \text{logit}^{-1}\left(\beta_0^f + \beta_{trt}^f + \beta_z^f z + \beta_{z:trt}^f z\right) \\ b(z) &= e^{(\beta_0^b + \beta_{trt}^b + \beta_z^b z + \beta_{z:trt}^b z)} \\ r_r &= \beta_0^r \\ r_d &= \text{TruncNormal}_L^U(\mu^d, \sigma^d) \end{aligned} \quad (20)$$

*Integration rule*: midpoint

L = 0.518

U = 3.722

Number of meshpoints: 400

***Erigeron speciosus:*** The large number of capitula and seeds prevented us from feasibly counting every seed. We therefore split the seed production function of the generic vital rate model into a function for the number of capitulae produced and the number of seeds per capitulum (Eq. 21).

$$\begin{aligned} &\text{Number of capitulae} \begin{cases} \left[Y_i^c | Y_i^f = 1\right] & \sim \text{GammaPoisson}(p_i^c, \theta_i^c) \\ \log(p_i^c) & = \beta_0^c + \beta_{trt}^c + \beta_z^c z_i + \beta_{z:trt}^c z_i + \alpha_{plot[j]}^c + \alpha_{year[k]}^c \\ \log(\theta_i^c) & = \gamma_0^c + \gamma_{trt}^c + \tau_{plot}^c + \tau_{year}^c \end{cases} \\ &\text{Seeds per capitulum} \begin{cases} \left[Y_i^p | Y_i^f = 1\right] & \sim \text{HurdleGamma}\left(\pi_i^p, \alpha_i^p, \beta_i^p\right) \\ \text{logit}\left(\pi_i^p\right) & = \beta_0^p + \beta_{trt}^p + \alpha_{plot[j]}^p + \alpha_{year[k]}^p \\ \log\left(\alpha_i^p\right) & = \gamma_0^p + \gamma_{trt}^g + \gamma_z^p z_i + \gamma_{z:trt}^p z_i + \tau_{plot[j]}^p + \tau_{year[k]}^p \\ \log\left(\beta_i^p\right) & = \delta_0^p + \delta_{trt}^p + \delta_z^p z_i + \delta_{z:trt}^p z_i + \omega_{plot[j]}^p + \omega_{year[k]}^p \end{cases} \end{aligned} \quad (21)$$

The $\alpha$, $\tau$, and $\omega$ varying effects for plot and year were included in the correlated varying effects (Eq. 5).

*IPM kernel definitions*:

$$\boldsymbol{P(z', z)}$$

$$\begin{aligned} s(z) &= \text{logit}^{-1}(\beta_0^s + \beta_{trt}^s + \beta_z^s z_i + \beta_{z:trt}^s z_i) \\ g(z', z) &= \text{TruncNormal}_L^U(\mu^g, \sigma^g) \\ \mu^g &= \beta_0^g + \beta_{trt}^g + \beta_z^g z + \beta_{z:trt}^g z \\ \sigma^g &= \gamma_0^g + \gamma_{trt}^g \end{aligned} \quad (22)$$

$$\boldsymbol{F(z', z)}$$

$$\begin{aligned} f(z) &= \text{logit}^{-1}\left(\beta_0^f + \beta_{trt}^f + \beta_z^f z + \beta_{z:trt}^f z\right) \\ b(z) &= e^{(\beta_0^b + \beta_{trt}^b + \beta_z^b z + \beta_{z:trt}^b z)} \\ r_r &= \beta_0^r \\ r_d &= \text{TruncNormal}_L^U(\mu^d, \sigma^d) \end{aligned} \quad (23)$$

*Integration rule*: midpoint

L = -2.632

U = 10.349

Number of meshpoints: 400

*Accounting for correlated spatiotemporal impacts on demographic vital rates*

The vital rate models for each species included correlations between group-level varying intercepts for plot and for year to account for spatiotemporal variation that impacted plant allocation constraints. We used $\text{LKJ}(10)$ as the prior distributions for the intercept correlation matrix of each group. This prior is skeptical of strong correlations, with >99% of the probability density concentrated between ±0.5.

None of the four plant species studied—*Delphinium nuttallianum*, *Hydrophyllum fendleri*, *Potentilla pulcherrima*, or *Erigeron speciosus*—had detectable correlations between the vital rate parameters at the plot or year levels (Figs. S6–S9). Moreover, the posterior distributions did not differ substantially from the prior distribution.

*Life Table Response Experiment contributions of individual parameters*

The Life Table Response Experiment (LTRE) calculated the contributions of each vital rate parameter to the treatment effects on λ. We summed these lower-level vital rate contributions to determine the net contributions of vital rate functions (Fig. 2, main text). Here, we present the parameter-level LTRE decomposition for each of the four plant species we studied (Figs. S10–S13). The LTRE contributions of parameters within the same vital rate function tended to be strongly correlated (Figs. S14–S17), meaning that the uncertainty around the vital rate function-level summed contributions may be amplified or attenuated compared to the uncertainty around the parameter-level contributions.

***2f. Elasticity partitioning***

The elasticity of $\lambda$ to element-wise perturbations of a projection matrix **A** is defined by de Kroon et al. (1986) as,

$$e_{ij} = \frac{\partial \ln \lambda}{\partial \ln a_{ij}} = \frac{a_{ij}}{\lambda}\frac{\partial \lambda}{\partial a_{ij}}, \qquad (24)$$

and the sum across the matrix of elasticities is,

$$\sum_{i,j} e_{ij} = 1. \qquad (25)$$

This sum-to-one constraint of the elasticity matrix allows for different biologically meaningful partitions to be quantified by summing subsets of the $e_{ij}$ elements.

Our goal was to decompose the elasticity matrix into two components that correspond to persistence (*sensu lato* to include established plants and seeds in the seed bank) and reproduction. The discretized IPM projection kernel, **A**, is a $2 \times 2$ block matrix (or "megamatrix") with elements $a_{ij}$,

$$\mathbf{A} = \begin{bmatrix} \mathbf{P} + \mathbf{F} & \mathbf{B}_{\text{out}} \\ \mathbf{B}_{\text{in}} & \mathbf{B}_{\text{stay}} \end{bmatrix}. \qquad (26)$$

The blocks contain discretized subkernels for survival and growth of established plants ($\mathbf{P}$), reproduction through immediately germinating seeds ($\mathbf{F}$), reproduction through seeds entering the seed bank ($\mathbf{B}_{\text{in}}$), seed bank survival ($\mathbf{B}_{\text{stay}}$), and germination from the seed bank ($\mathbf{B}_{\text{out}}$). The blocks can be partitioned into the desired life cycle components as,

$$\mathbf{A} = \underbrace{\begin{bmatrix} \mathbf{P} & 0 \\ 0 & \mathbf{B}_{\text{stay}} \end{bmatrix}}_{\text{persistence}} + \underbrace{\begin{bmatrix} \mathbf{F} & \mathbf{B}_{\text{out}} \\ \mathbf{B}_{\text{in}} & 0 \end{bmatrix}}_{\text{reproduction}}. \qquad (27)$$

Seed survival in the seed bank is included in the persistence partition, whereas the seeds going into or germinating from the seed bank are included in the reproduction partition. In the absence of a seed bank, these partitions reduce to a simple matrix $\mathbf{A} = \mathbf{P} + \mathbf{F}$.

In the first block of the Equation 27 partitions, the elements of $\mathbf{A}$ are the sum of contributions from the persistence and reproduction partitions, $a_{ij} = p_{ij} + f_{ij}$, that must be separated. For each of these elements $a_{ij}$, we calculated the proportion of the transition probability attributable to the persistence and reproductive partitions as $\pi_{ij} = p_{ij}/a_{ij}$ and $\phi_{ij} = f_{ij}/a_{ij}$, respectively. In the other three blocks, the value of $a_{ij}$ comes from a single partition, reducing these proportions to either 0 or 1 while the other is 1 or 0. Weighting each elasticity by the proportional contribution of each partition gives the decomposition of Equation 25 as,

$$\begin{aligned} \sum_{i,j} e_{ij} &= \sum_{i,j} e_{ij} \left(\pi_{ij} + \phi_{ij}\right) \\ &= \underbrace{\sum_{i,j} e_{ij}\, \pi_{ij}}_{\text{persistence}} + \underbrace{\sum_{i,j} e_{ij}\, \phi_{ij}}_{\text{reproduction}} \qquad (28) \\ &= 1. \end{aligned}$$

Thus the elasticities of $\lambda$ to element-wise perturbations of $\mathbf{A}$ can be exactly partitioned into persistence and reproduction components of the life cycle for the structure of our IPM. Moreover, these are partitions of the unit simplex which allows comparisons both within and among species.

**Table S1.** Percent change in both flower- and plant-level seed production are poor predictors of population growth rate, λ. Values are calculated from raw data: means for each metric across all reproductive individuals and all four years of the experiment. For *H. fendleri* and *E. speciosus*, data are seeds per inflorescence and per capitula or flowerhead, respectively, as each inflorescence and capitula contain numerous flowers and were not counted due to time constraints.

| **Species** | **Treatment comparison** | **% Change seeds per flower** | **% Change seeds per plant** | **% Change λ** |
|---|---|---|---|---|
| *D. nuttallianum* | control vs. reduced | -63.4 | -58.0 | -8.7 |
| | control vs. increased | +2.8 | +4.7 | -5.9 |
| *H. fendleri* | control vs. reduced | -52.6 | -62.3 | +0.8 |
| | control vs. increased | +11.8 | -7.3 | -3.0 |
| *P. pulcherrima* | control vs. reduced | -15.3 | -25.7 | +2.1 |
| | control vs. increased | +7.3 | -0.7 | -3.4 |
| *E. speciosus* | control vs. reduced | -56.6 | -40.4 | +4.1 |
| | control vs. increased | +17.7 | +19.2 | +9.7 |

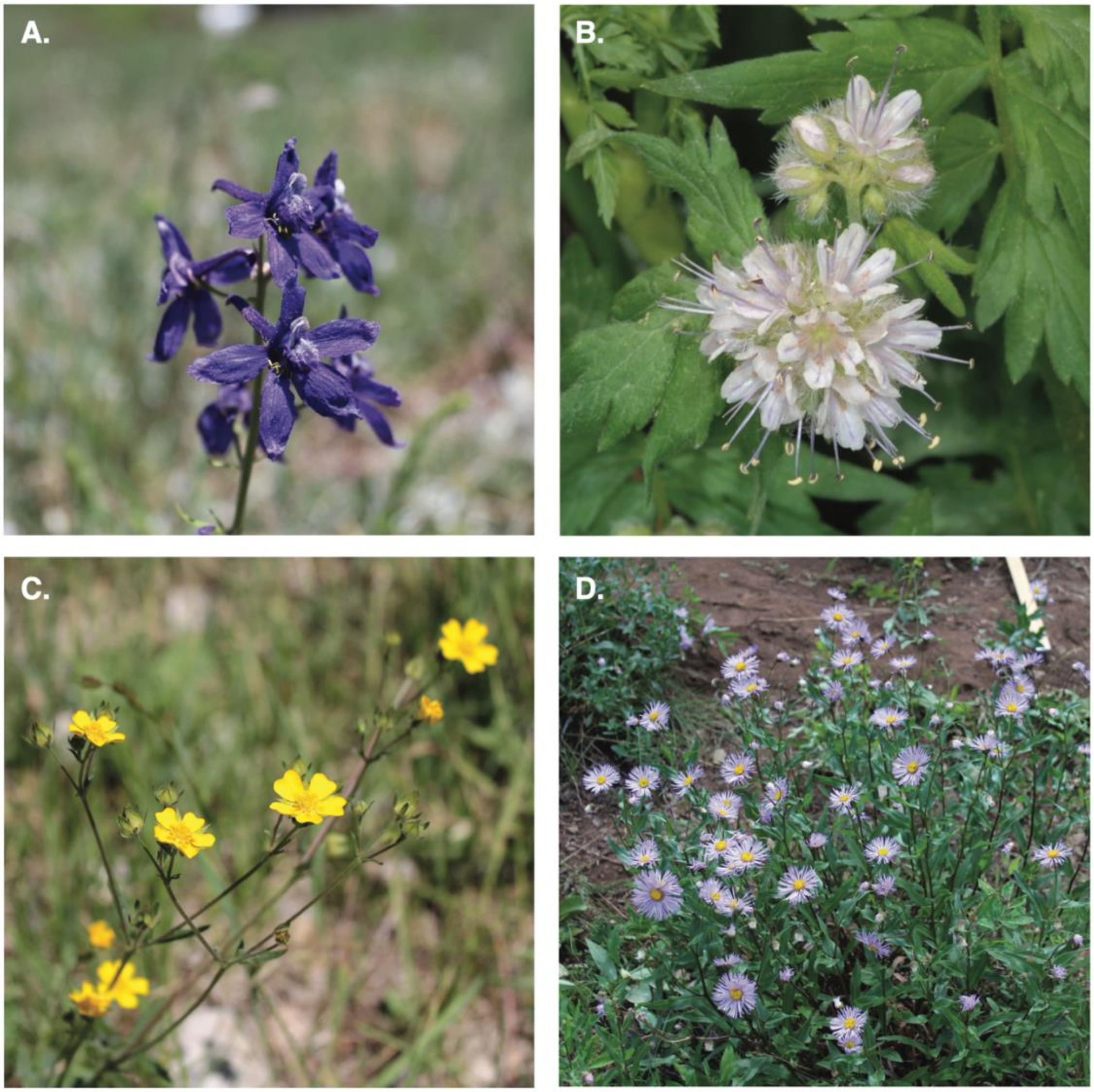


**Fig. S3.** Photographs of the four study species. (A) *Delphinium nuttallianum* (Ranunculaceae), (B) *Hydrophyllum fendleri* (Hydrophyllaceae), (C) *Potentilla pulcherrima* (Rosaceae), and (D) *Erigeron speciosus* (Asteraceae). Photographs in A, C, and D by P.J. CaraDonna, and in B by D.W. Inouye.

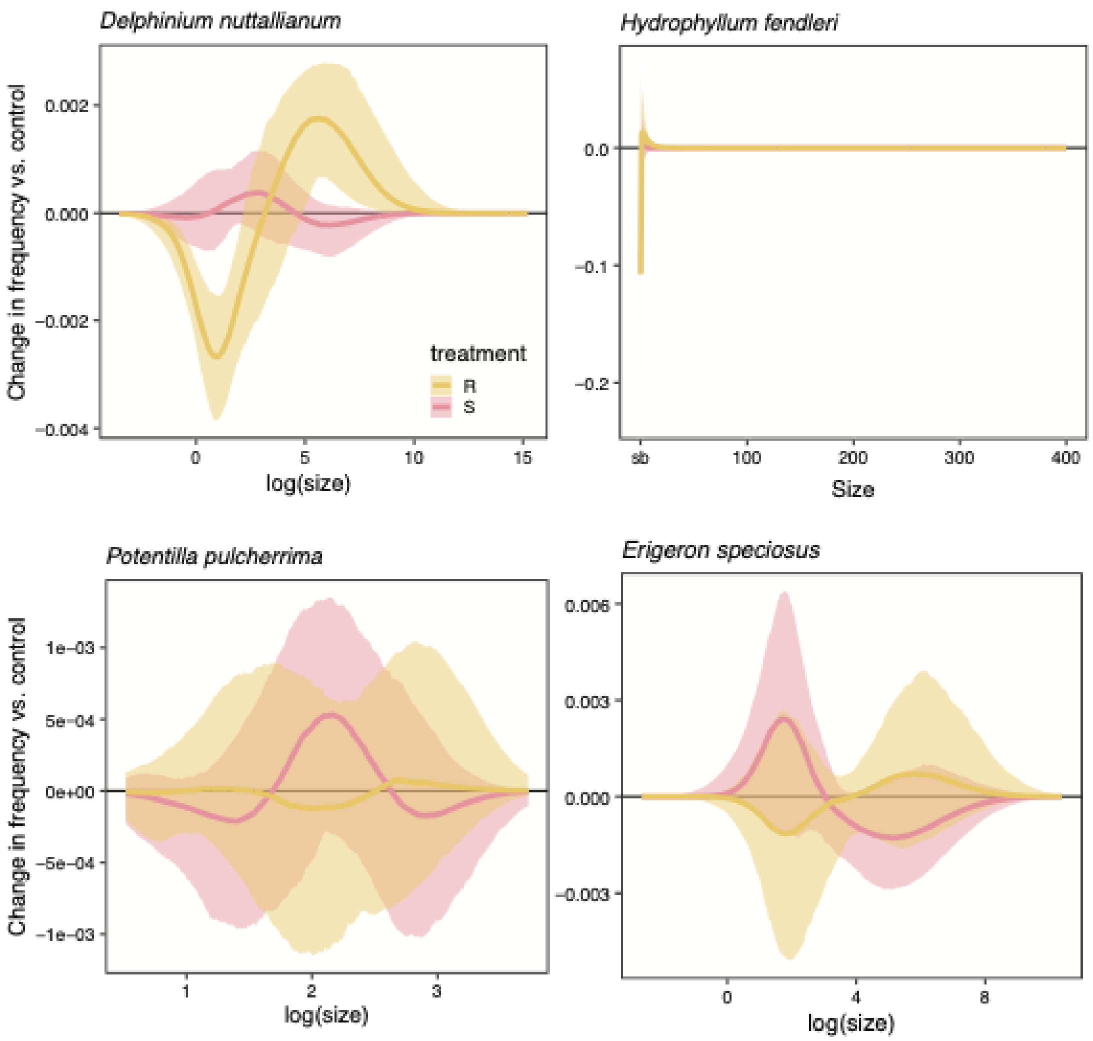


**Fig. S4.** Stable size structure of each species, shown relative to the control (0 on the y-axis). R stands for reduced pollination (decreased pollination treatment) and S for supplemental pollination (increased pollination treatment). For all species, treatments overlap with the control, except *D. nuttallianum* in the reduced pollination treatment. Even so, areas where it differs are small in magnitude, showing that our treatments did not fundamentally change the size structure of our study populations.

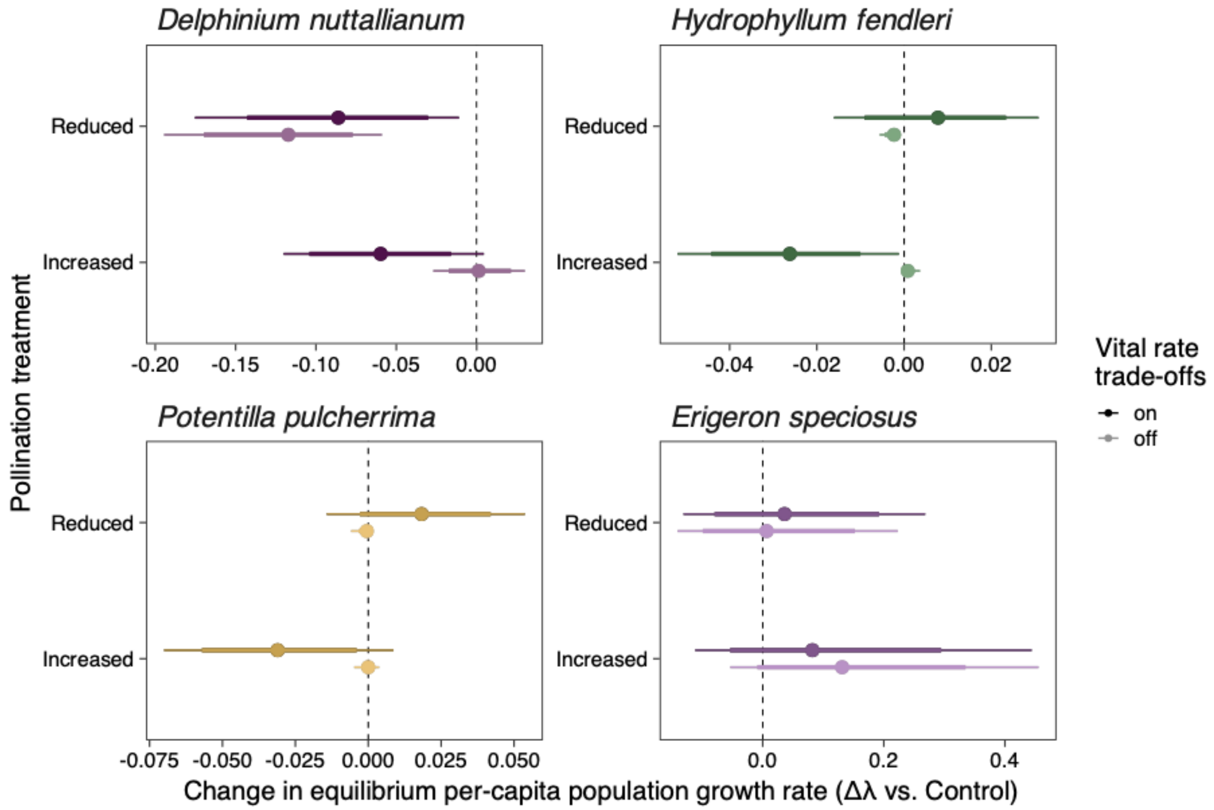


**Fig. S5.** Vital rate trade-offs are important for understanding how population growth rate, $\lambda$, responds to pollination treatments. Posterior distribution summaries of the change in population growth rate, $\lambda$, under increased and decreased pollination treatments relative to the control, with and without demographic vital rate trade-offs. Trade-offs were turned ‘off’ by limiting the pollination treatment effects to seed production and substituting parameters estimated from the controls for all other vital rates. For cases when pollination treatments affect $\lambda$ in unexpected directions (*e.g.*, decrease in $\lambda$ in response to increased pollination), turning trade-offs ‘off’ results in little to no change in $\lambda$ relative to controls. For cases when pollination treatments affect $\lambda$ in expected directions (*D. nuttallianum* reduced vs. control and *E. speciosus* increased vs. control), effects of treatments are magnified when trade-offs are turned ‘off’, because there is nothing to counteract the change in seed production. Points show the posterior median and line ranges show the 75% (thick line) and 90% (thin line) credible intervals.

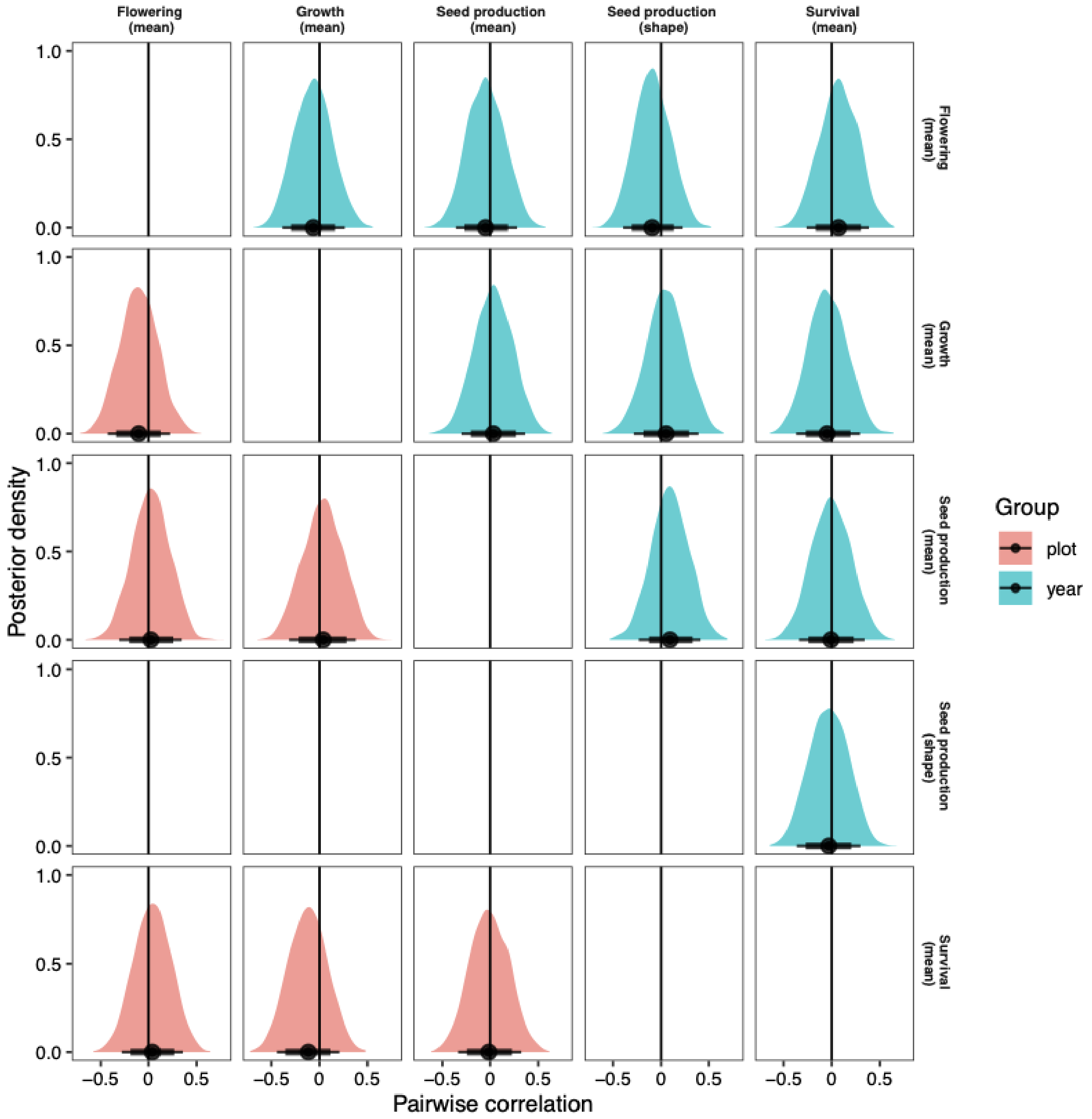


**Fig. S6.** There were no detectable correlations between the vital rate parameters at the plot or year levels in the joint vital rate model fit for *Delphinium nuttallianum*. Each panel shows the posterior distribution of the pairwise correlation coefficients between group-level intercepts for plot (red) and year (blue). The row and column labels of each panel report the vital rate function component, and the correlated intercepts inform the parameter of the response distribution indicated in parentheses (usually the mean/location, but also other distributional parameters depending on the vital rate component). Point summaries show the median, and the whiskers show the 75% (thick line) and 90% (thin line) credible intervals.

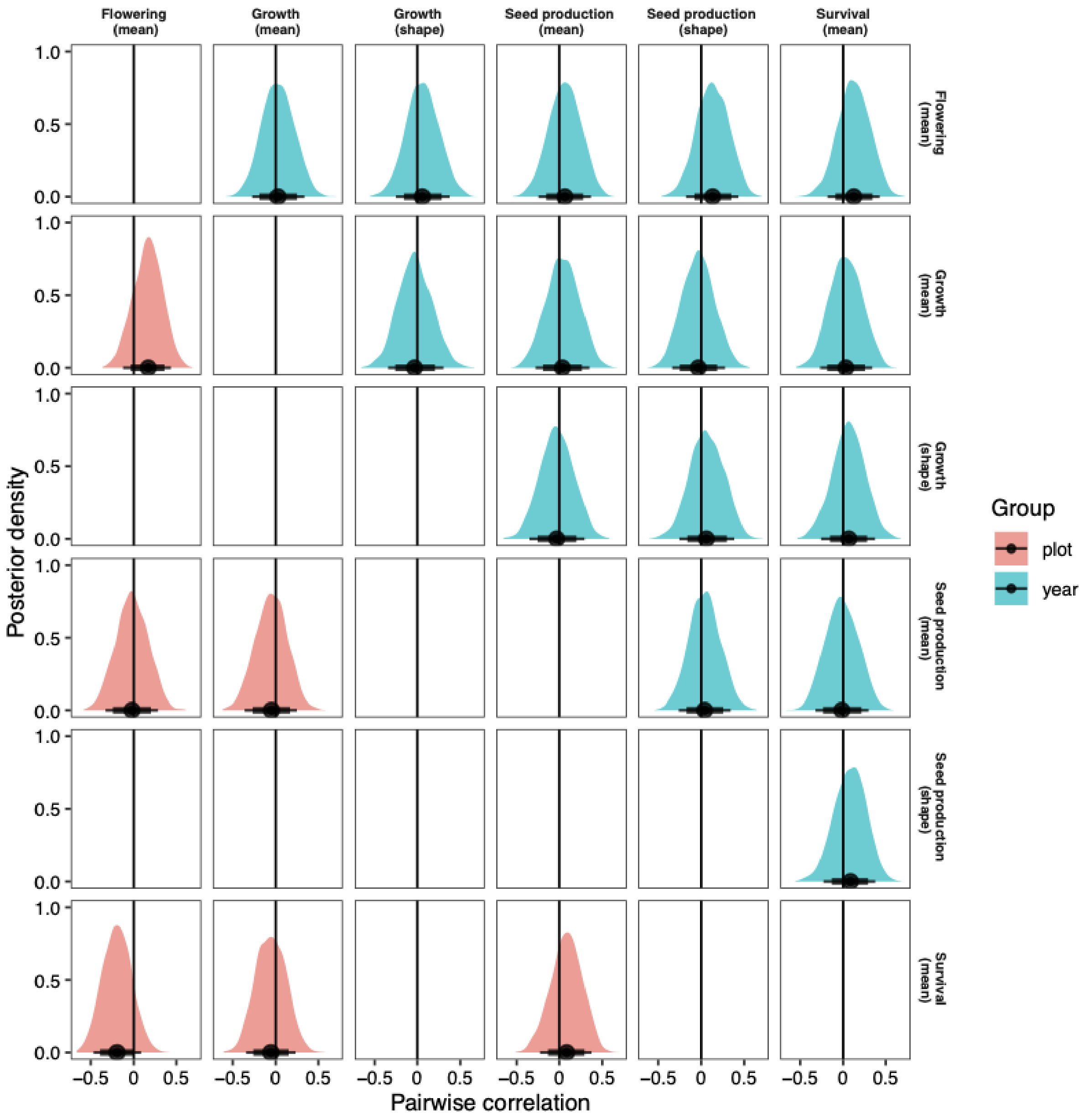


**Fig. S7.** There were no detectable correlations between the vital rate parameters at the plot or year levels in the joint vital rate model fit for *Hydrophyllum fendleri*. Each panel shows the posterior distribution of the pairwise correlation coefficients between group-level intercepts for plot (red) and year (blue). The row and column labels of each panel report the vital rate function component, and the correlated intercepts inform the parameter of the response distribution indicated in parentheses (usually the mean/location, but also other distributional parameters depending on the vital rate component). Point summaries show the median, and the whiskers show the 75% (thick line) and 90% (thin line) credible intervals.

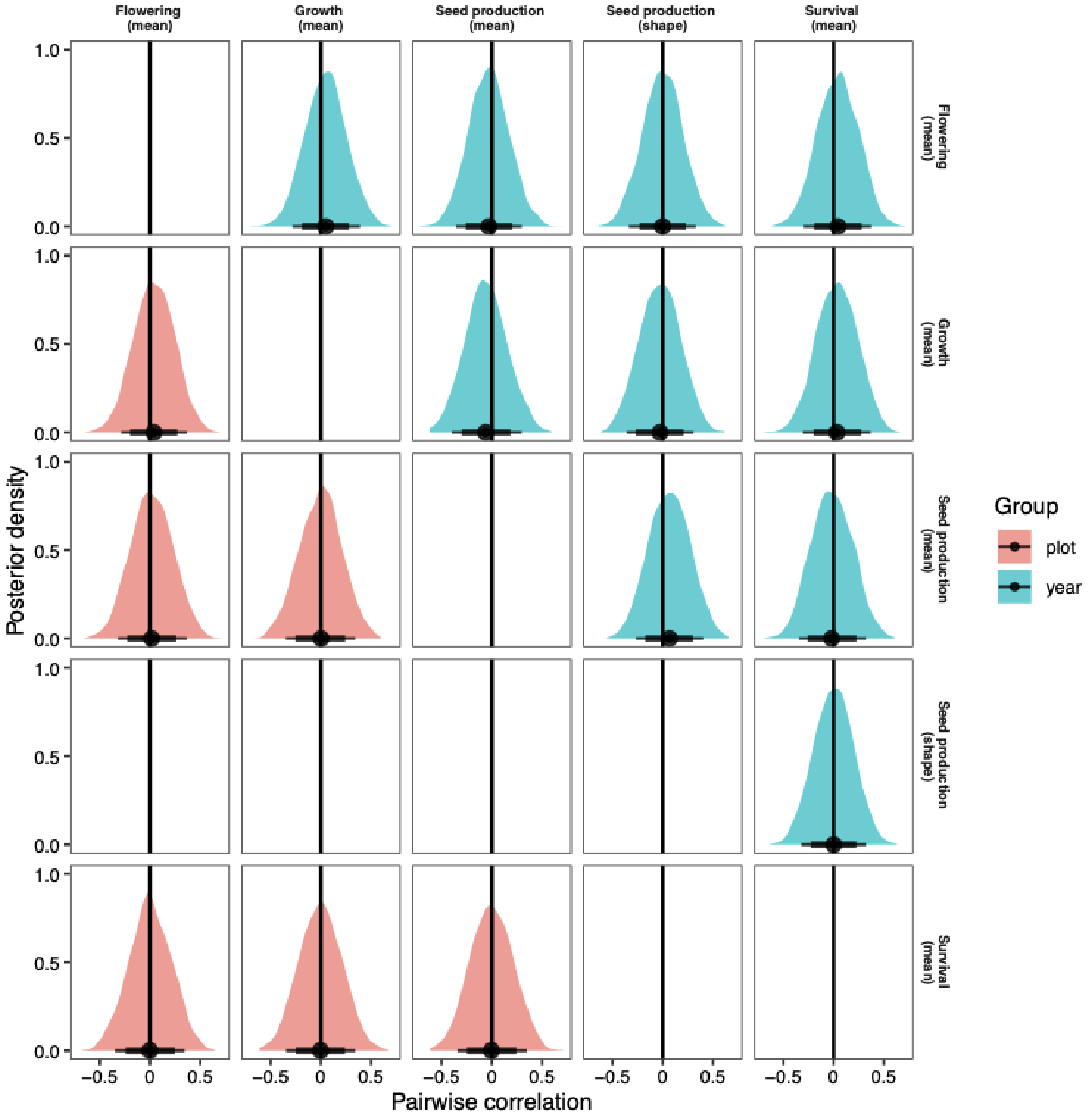


**Fig. S8.** There were no detectable correlations between the vital rate parameters at the plot or year levels in the joint vital rate model fit for *Potentilla pulcherrima*. Each panel shows the posterior distribution of the pairwise correlation coefficients between group-level intercepts for plot (red) and year (blue). The row and column labels of each panel report the vital rate function component, and the correlated intercepts inform the parameter of the response distribution indicated in parentheses (usually the mean/location, but also other distributional parameters depending on the vital rate component). Point summaries show the median, and the whiskers show the 75% (thick line) and 90% (thin line) credible intervals.

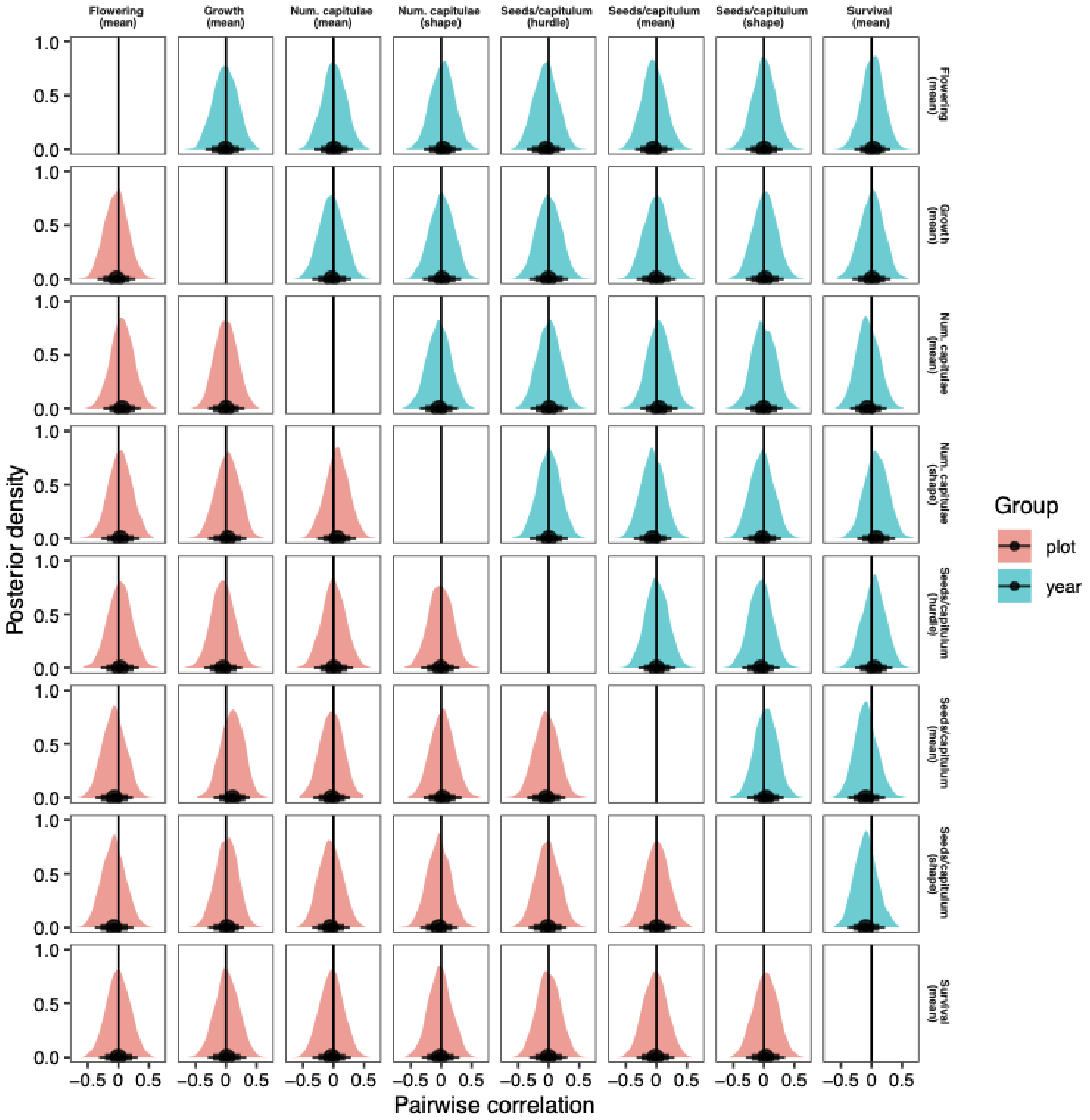


**Fig. S9.** There were no detectable correlations between the vital rate parameters at the plot or year levels in the joint vital rate model fit for *Erigeron speciosus*. Each panel shows the posterior distribution of the pairwise correlation coefficients between group-level intercepts for plot (red) and year (blue). The row and column labels of each panel report the vital rate function component, and the correlated intercepts inform the parameter of the response distribution indicated in parentheses (usually the mean/location, but also other distributional parameters depending on the vital rate component). Point summaries show the median, and the whiskers show the 75% (thick line) and 90% (thin line) credible intervals.

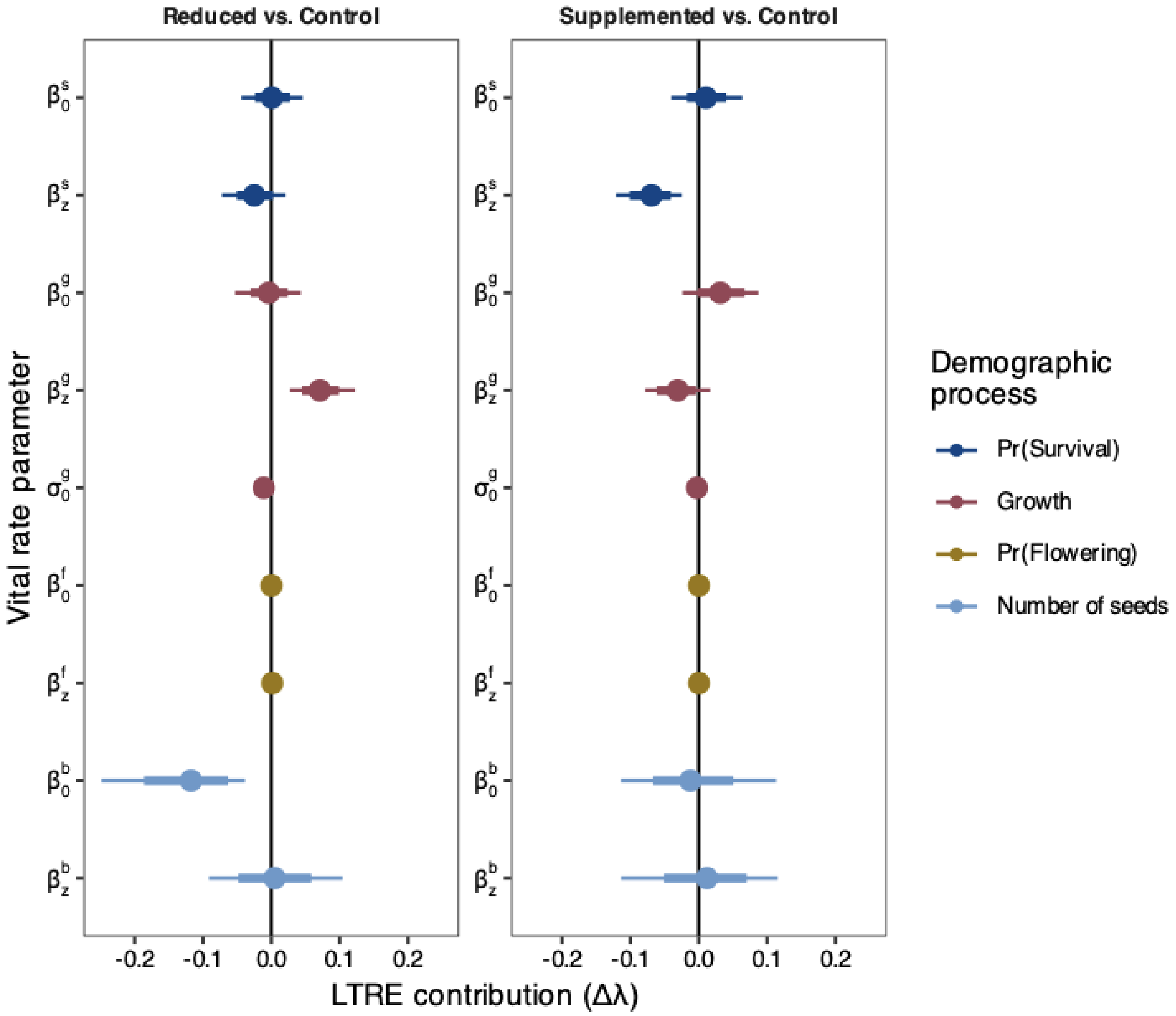


**Fig. S10.** Parameter-level Life Table Response Experiment contributions for *Delphinium nuttallianum*. Points are the median of the posterior distribution, and the whiskers show the 67% (thick) and 90% (thin) credible intervals.

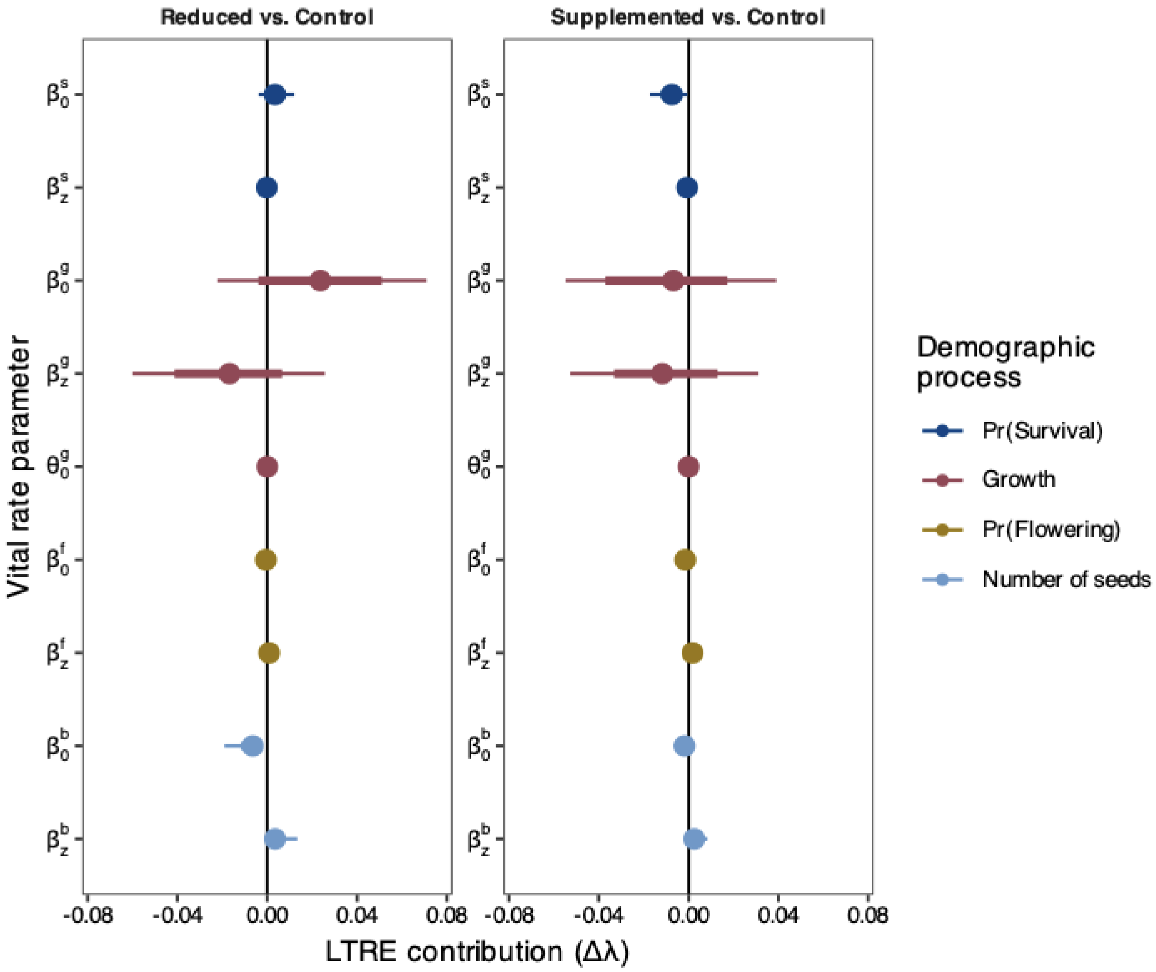


**Fig. S11.** Parameter-level Life Table Response Experiment contributions for *Hydrophyllum fendleri*. Points are the median of the posterior distribution, and the whiskers show the 67% (thick) and 90% (thin) credible intervals.

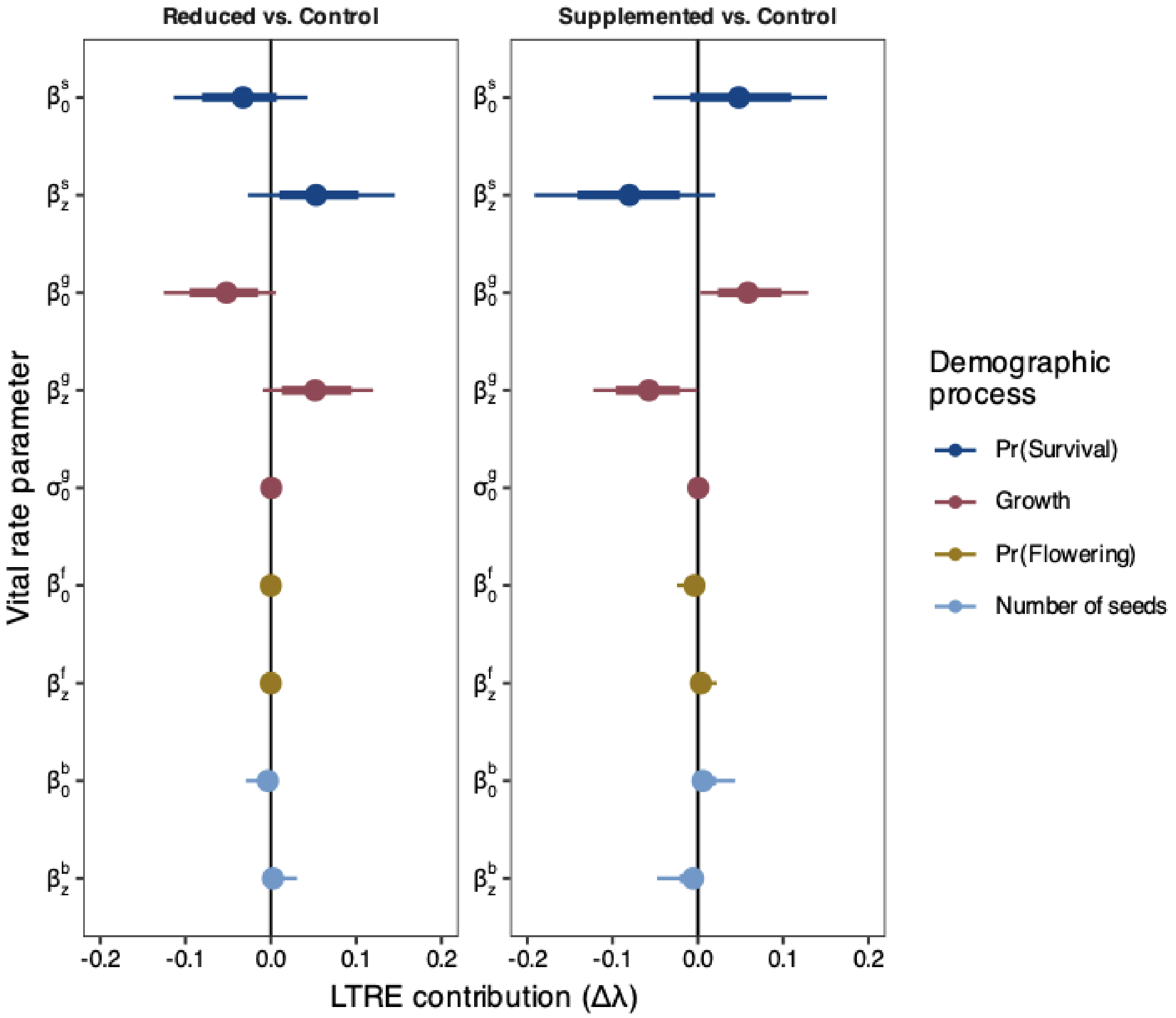


**Fig. S12.** Parameter-level Life Table Response Experiment contributions for *Potentilla pulcherrima*. Points are the median of the posterior distribution, and the whiskers show the 67% (thick) and 90% (thin) credible intervals.

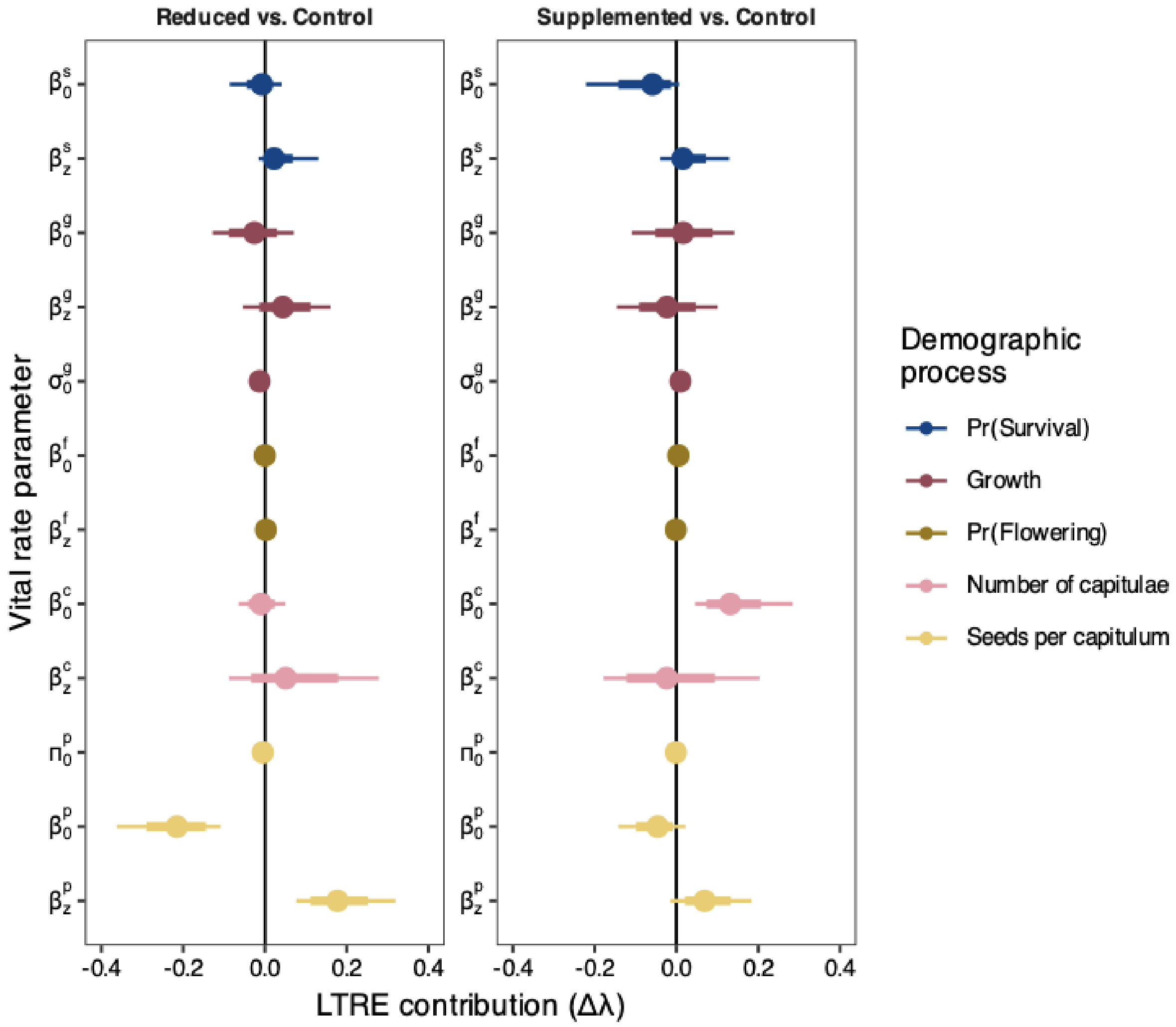


**Fig. S13.** Parameter-level Life Table Response Experiment contributions for *Erigeron speciosus*. Points are the median of the posterior distribution, and the whiskers show the 67% (thick) and 90% (thin) credible intervals.

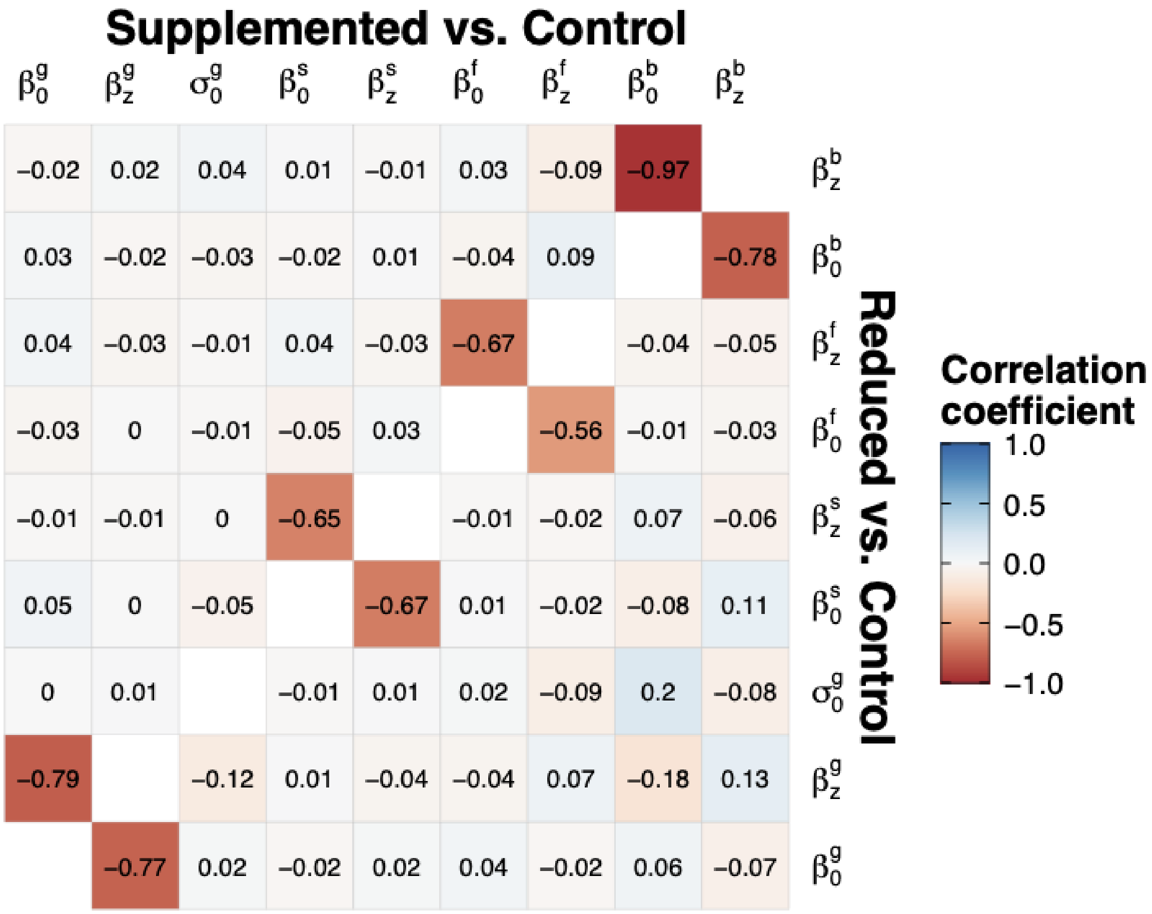


**Fig. S14.** Posterior correlations between the parameter-level Life Table Response Experiment contributions in *Delphinium nuttallianum*. The upper triangle shows the Supplemented vs. Control contrast, and the lower triangle shows the Reduced vs. Control contrast. Parameter definitions are in Eq. 4, with $\beta_0$ representing intercepts and $\beta_z$ slopes from each vital rate model. $\sigma_0^g$ represents the standard deviation of the growth function. Superscripts g = growth, s = survival, f = probability of flowering, and b = number of seeds.

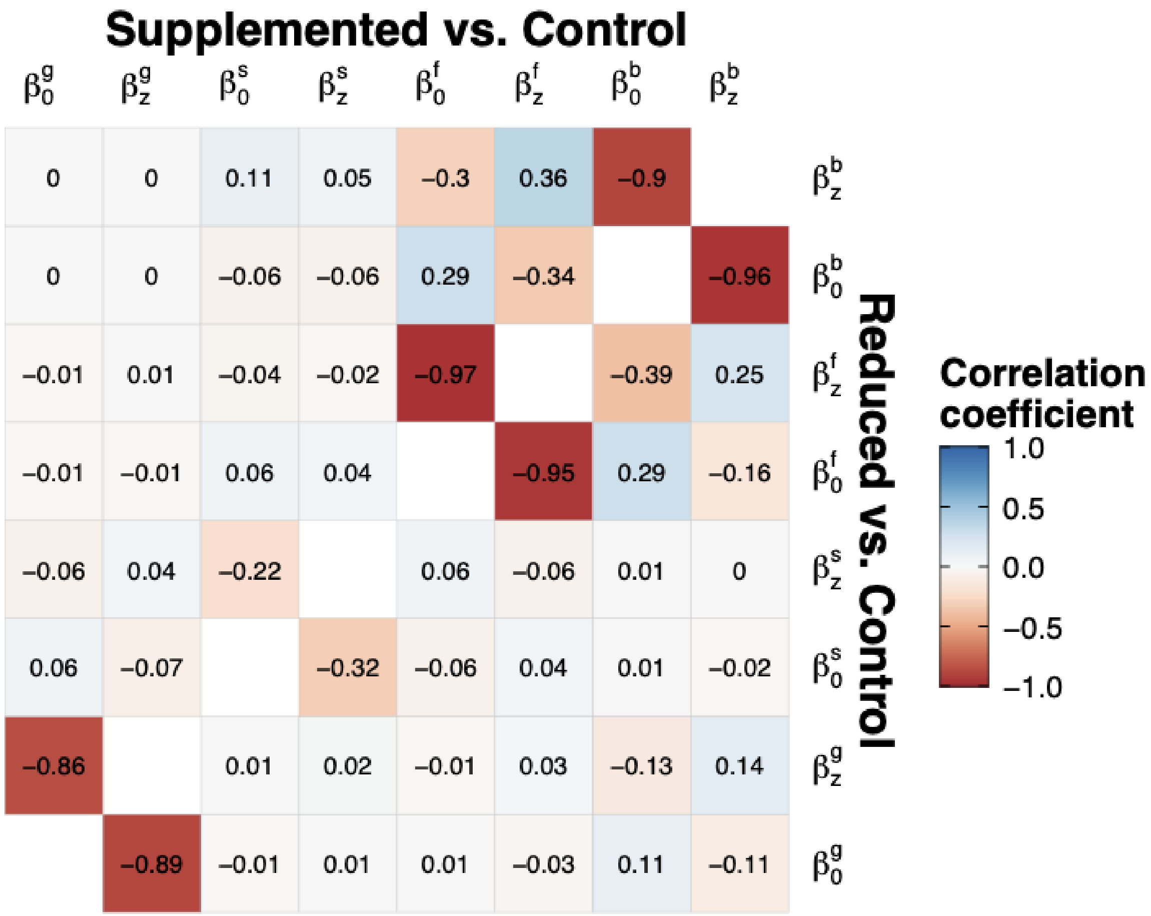


**Fig. S15.** Posterior correlations between the parameter-level Life Table Response Experiment contributions in *Hydrophyllum fendleri*. The upper triangle shows the Supplemented vs. Control contrast, and the lower triangle shows the Reduced vs. Control contrast. Parameter definitions are in Eq. 4, with $\beta_0$ representing intercepts and $\beta_z$ slopes from each vital rate model. $\sigma_0^g$ represents the standard deviation of the growth function. Superscripts g = growth, s = survival, f = probability of flowering, and b = number of seeds.

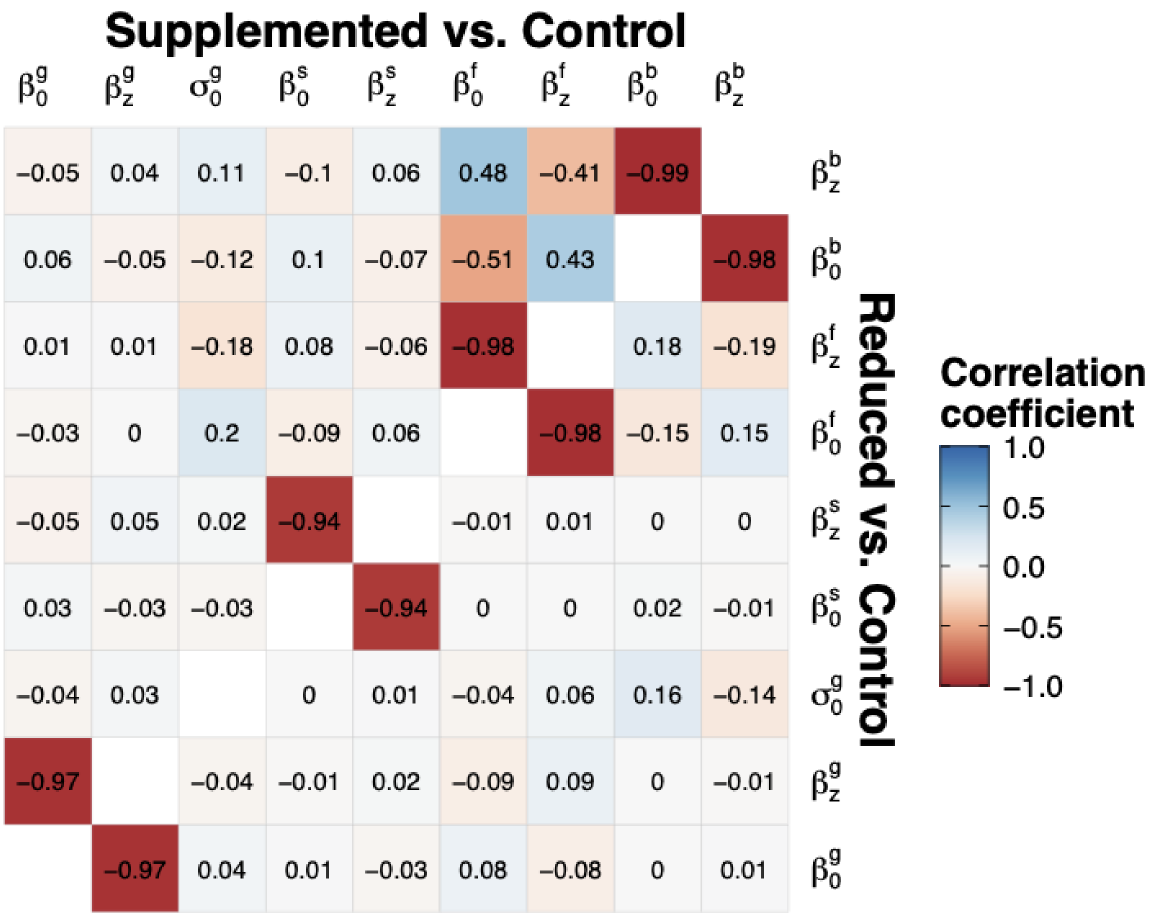


**Fig. S16.** Posterior correlations between the parameter-level Life Table Response Experiment contributions in *Potentilla pulcherrima*. The upper triangle shows the Supplemented vs. Control contrast, and the lower triangle shows the Reduced vs. Control contrast. Parameter definitions are in Eq. 4, with $ß_0$ representing intercepts and $ß_z$ slopes from each vital rate model. $\sigma_0^g$ represents the standard deviation of the growth function. Superscripts g = growth, s = survival, f = probability of flowering, and b = number of seeds.

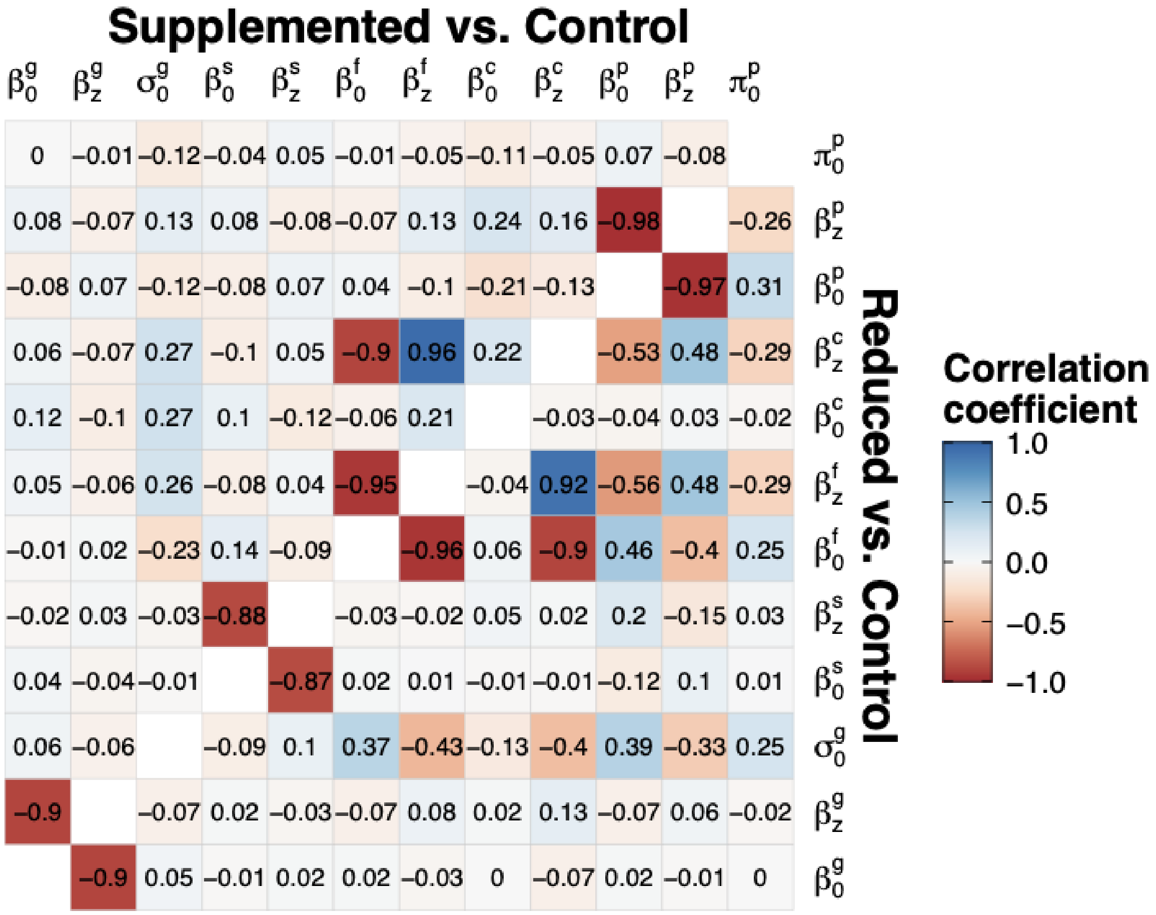


**Fig. S17.** Posterior correlations between the parameter-level Life Table Response Experiment contributions in *Erigeron speciosus*. The upper triangle shows the Supplemented vs. Control contrast, and the lower triangle shows the Reduced vs. Control contrast. Parameter definitions are in Eq. 4, with $\beta_0$ representing intercepts and $\beta_z$ slopes from each vital rate model. $\sigma_0^g$ represents the standard deviation of the growth function. Superscripts g = growth, s = survival, f = probability of flowering, and c = number of capitulae, p = seeds per capitulum.
